# LEM2 recruits CHMP7 for ESCRT-mediated nuclear envelope closure in fission yeast and human cells

**DOI:** 10.1101/049312

**Authors:** Mingyu Gu, Dollie LaJoie, Opal S. Chen, Alexander Von Appen, Mark S. Ladinsky, Michael J. Redd, Linda Nikolova, Pamela J. Bjorkman, Wesley I. Sundquist, Katharine S. Ullman, Adam Frost

## Abstract

ESCRT-III proteins have been implicated in sealing the nuclear envelope in mammals, spindle pole body dynamics in fission yeast, and surveillance of defective nuclear pore complexes in budding yeast. Here, we report that Lem2p (LEM2), a member of the LEM (Lap2-Emerin-Man1) family of inner nuclear membrane proteins, and the ESCRT-II/ESCRT-III hybrid protein Cmp7p (CHMP7), work together to recruit additional ESCRT-III proteins to holes in the nuclear membrane. In *S. pombe*, deletion of the ATPase *vps4* leads to severe defects in nuclear morphology and integrity. These phenotypes are suppressed by loss-of-function mutations that arise spontaneously in *lem2* or *cmp7,* implying that these proteins may function upstream in the same pathway. Building on these genetic interactions, we explored the role of LEM2 during nuclear envelope reformation in human cells. We found that CHMP7 and LEM2 enrich at the same region of the chromatin disc periphery during this window of cell division, and that CHMP7 can bind directly to the C-terminal domain of LEM2 *in vitro*. We further found that, during nuclear envelope formation, recruitment of the ESCRT factors CHMP7, CHMP2A and IST1/CHMP8 all depend on LEM2 in human cells. We conclude that Lem2p/LEM2 is a conserved nuclear site-specific adaptor that recruits Cmp7p/CHMP7 and downstream ESCRT factors to the nuclear envelope.

## Significance

The molecular mechanism for sealing newly formed nuclear envelopes was unclear until the recent discovery that ESCRT-III proteins mediate this process. CHMP7, in particular, was identified as an early-acting factor that recruits other ESCRT-III proteins to the nuclear envelope. A fundamental aspect of the varied roles of ESCRT factors is their recruitment by site-specific adaptors, yet the central question of how the ESCRT machinery is targeted to nuclear membranes has remained outstanding. Our study identifies the inner nuclear membrane protein LEM2 as a key, conserved factor that recruits CHMP7 and downstream ESCRT-III proteins to breaches in the nuclear envelope.

## Introduction

Eukaryotic genomes are secluded within the nucleus, an organelle with a boundary that comprises the double-membraned nuclear envelope (1). The inner and outer bilayers of the nuclear envelope are perforated by annular channels that contain Nuclear Pore Complexes (NPCs), each a massive assembly that regulates the trafficking of macromolecules like mRNA and proteins between the cytoplasm and nucleoplasm. The evolution of the envelope, among other roles, helped safeguard genome duplication and mRNA transcription from parasitic nucleic acids (2). The isolation of nucleoplasm from cytoplasm, however, presents a challenge during cell division when duplicated chromosomes must be separated for daughter cell inheritance.

Chromosome inheritance depends on assembly of a mitotic spindle, which pulls chromosomes toward opposite sides of the duplicating cell. Spindle assembly begins when two microtubule organizing centers (MTOCs) nucleate polymerization of anti-parallel arrays of microtubules in order to capture daughter chromosomes. Despite functional conservation throughout Eukarya, the mechanisms by which spindle microtubules breach the nuclear envelope to gain access to metaphase chromosomes vary markedly (3-6). In vertebrates and other organisms that have an “open mitosis”, the nuclear envelope disassembles completely, so that nucleoplasmic identity is lost. Certain protists and fungi, by contrast, maintain nuclear envelope integrity throughout a “closed mitosis” (3, 5).

The fission yeast *Schizosaccharomyces pombe* and the budding yeast *Saccharomyces cerevisiae* integrate their MTOC, known as the spindle pole body (SPB), into the nuclear envelope so that microtubule assembly can occur in the nucleoplasm (3, 4, 7, 8). In budding yeast, duplication of the SPB is coupled with nuclear envelope remodeling so that mother and daughter SPBs reside within the envelope throughout the cell cycle (4, 9). Fission yeast, by contrast, restrict SPB access to the nucleoplasm during mitosis only (7, 8). Upon mitotic entry, a fenestration through the nuclear envelope opens transiently and the mother and the daughter SPBs are incorporated (3, 5, 6, 8, 10). For every cell cycle, therefore, the fission yeast nuclear envelope must open and reseal twice: once when the SPBs are inserted, and again when the SPB is ejected from the envelope following a successful cell cycle.

Incorporating NPCs and SPBs into the nuclear envelope requires certain factors and mechanisms in common, including membrane remodeling activities (6, 11-15). We and others have previously reported strong genetic interactions between transmembrane nucleoporins, spindle pole body components, and endosomal sorting complexes required for transport (ESCRT) genes—portending a role for certain ESCRT proteins in nuclear membrane remodeling (16, 17). In general, ESCRT components are recruited to different target membranes by site-specific adaptors that ultimately recruit the membrane remodeling ESCRT-III subunits and their binding partners, including VPS4-family adenosine triphosphatases (ATPases) (18-20). We previously showed that certain ESCRT-III mutants and *vps4Δ* cells displayed an apparent over-amplification of SPBs (or defective fragments) in fission yeast and that the severity of this SPB phenotype in fission yeast waned over time, suggesting possible genetic suppression (16). In budding yeast, Webster et al. reported that without ESCRT-III/Vps4 activity, misassembled NPCs accumulate in a compartment they named the SINC for Storage of Improperly Assembled Nuclear Pore Complexes (21). They also showed that LEM family (Lap2-Emerin-Man1) inner nuclear membrane proteins, Heh1p and Heh2p in budding yeast, associate with defective NPC assembly intermediates (but not with mature NPCs), and that Heh1/2 proteins may recruit ESCRT–III and Vps4 activities to malformed NPCs in order to clear them from the nuclear envelope (21).

In mammals, VPS4 depletion induces nuclear morphology defects (22), and several recent reports have demonstrated that ESCRT pathway proteins are recruited transiently to seal gaps in re-forming mammalian nuclear membranes during anaphase (23, 24), and to rupture sites in the nuclei of interphase mammalian cells (25, 26). Depletion of ESCRT factors delays sealing of the reforming nuclear envelope and impairs mitotic spindle disassembly (23, 24). Moreover, depletion of SPASTIN, another meiotic clade VPS4-family member and ESCRT-III-binding enzyme (27), also delays spindle disassembly and envelope resealing (24, 28). Similar effects were seen upon depletion of several ESCRT-III proteins, including the poorly characterized ESCRT factor, CHMP7, which has features of both ESCRT-II and ESCRT-III proteins (29). These observations support a model in which ESCRT-III proteins and SPASTIN together coordinate microtubule severing with the closure of annular gaps in the nuclear envelope. This model is conceptually similar to the mechanism of cytokinetic abscission, where SPASTIN disassembles the residual microtubules that pass between daughter cells, while ESCRT-III and VPS4 proteins constrict the midbody membrane to the point of fission (19, 28, 30).

Here we address the key question of what upstream factor(s) serves as the membrane-specific adaptor that facilitates CHMP7 recruitment to function in sealing nuclear envelope breaches. To identify factors in this pathway, we returned to the genetically tractable fission yeast system. We report that deletion of *vps4* in *S. pombe* leads to severe defects in nuclear membrane morphology and nuclear integrity, with secondary defects in NPCs and SPB dynamics. Remarkably, these phenotypes are suppressed spontaneously when cells acquire loss-of-function mutations in *cmp7,* an ortholog of human CHMP7, or in *lem2,* a LEM domain inner nuclear membrane protein and ortholog of human LEM2. We also show that in human cells recruitment of CHMP7 and downstream ESCRT-III proteins to the re-forming nuclear envelope during anaphase depends on LEM2, most likely through a direct interaction between CHMP7 and the C-terminal nucleoplasmic domain of LEM2. Together, these observations implicate LEM2 as a nucleus-specific adaptor that recruits ESCRT pathway activities to remodel the nuclear envelope during both open and closed mitoses across Eukaryotes.

## Results

### *vps4∆* fission yeast cells grow very slowly and loss of either *cmp7* or *lem2* rescues growth

The AAA ATPase VPS4 has a primary role in disassembling ESCRT-III polymeric structures in the different settings where the ESCRT pathway mediates membrane remodeling. To determine whether and how previously-reported phenotypes that result from deletion of VPS4 were suppressed over time (16), we monitored the growth of individual colonies following sporulation and tetrad dissection of *vps4Δ/+* diploid cells. Growth rates of *vps4Δ* spores were dramatically slower than wild type spores (Fig 1A). This growth defect spontaneously reverted over time, so that when mutant spores were streaked on rich media, some *vps4Δ* colonies exhibited growth rates comparable to wild type colonies (Fig 1B). To identify potential suppressor mutations, we sequenced complete genomes for 12 strains that spontaneously reverted to wild type growth rates and compared them with genomes of both wild type and apparently unsuppressed *vps4Δ* strains. The analysis revealed that 7 out of the 12 suppressors had different loss-of-function mutations in the ESCRT-II/ESCRT-III hybrid gene, *cmp7* (Table S1). The remaining 5 each had equivalent independent mutations in a LEM domain family member, *lem2*, within what appears to be a slippery poly-T track (Table S1). These 12 mutant alleles were further confirmed by Sanger sequencing, and none of the suppressors were found to harbor mutations in both *cmp7* and *lem2* simultaneously.

**Figure 1.**
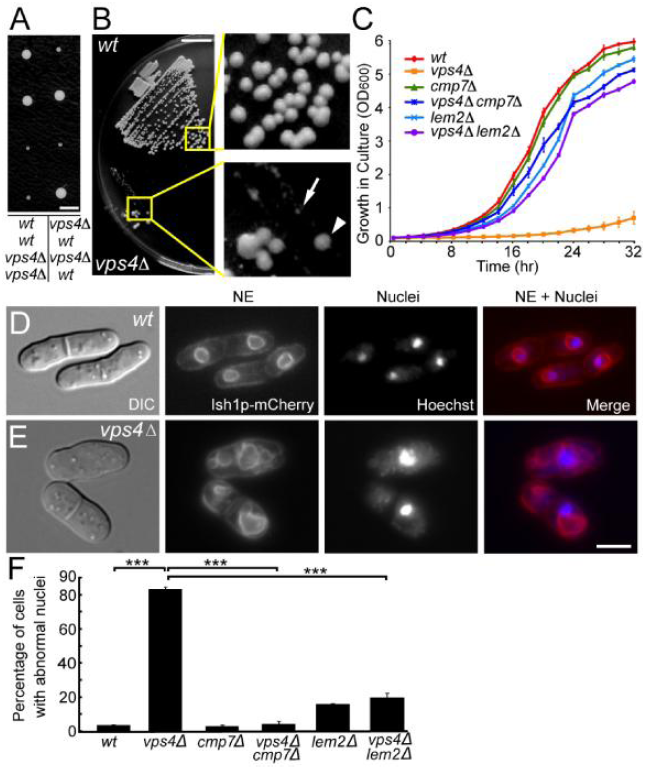
*vps4Δ* cells grow slowly and have severe nuclear envelope defects, which are suppressed by loss of *cmp7* or *lem2*. (A) Tetrad dissection of *vps4Δ/+* diploids, with genotypes labeled below the image. (Scale bar: 0.25 cm) (B) Spontaneous suppressors (arrowhead) of *vps4Δ* (arrow) appearing after three days on rich media, with colony sizes comparable to those of wild type (*wt*) cells. (Scale bar: 1 cm) (C) Growth curves of each genotype showing optical densities of yeast cultures, starting at OD600nm 0.06, for 32 hours in 2-hour intervals. Three independent isolates for each genotype were measured. The plots show mean±SEM. (D) *wt* cells, showing normal NE morphologies (Ish1p-mCherry) and DNA (Hoechst staining) within the nuclei. (E) *vps4Δ*, showing various NE morphology defects including excess NE, fragmented NE, and DNA that appears to be outside of the nucleus. (F) Deletion of either *cmp7* or *lem2* rescues these NE morphology defects. Here and throughout, three independent isolates/strains were imaged for each genotype, and n represents the total number of scored cells. Mean±SEM, *wild type*: 1±1%, n=115, 121, 139; *vps4Δ*: 85±5%, n=62, 69, 86; *cmp7Δ*: 2.0±0.4%, n=151, 166, 147; *vps4Δcmp7Δ*: 1.1±0.7%, n =161, 118, 145; *lem2Δ*: 8±3%, n= 131, 103, 128; *vps4Δlem2Δ*: 12.2±0.7%, n=99, 176, 111. Two-tail student *t* tests were used here and throughout and designated as follows: \**p*<0.05, \*\**p*<0.01, \*\*\**p*<0.001, N.S. “Not Significant”. (Scale bar: 5 μm).

To determine whether these potentially suppressive alleles rescued the growth of *vps4Δ* cells, we engineered *cmp*7*Δ/+* and *lem2Δ/+* genotypes within our *vps4Δ/+* diploid background and isolated *lem2Δ*, *cmp7Δ, vps4Δ* single mutants and their corresponding double mutants via sporulation and tetrad dissection. Quantitative growth rates in liquid culture for biological and technical triplicates demonstrated that *cmp7Δ* single mutants had wild-type growth rates and that the *vps4Δcmp7Δ* double mutant cells grew much faster than *vps4Δ* single mutant cells (albeit slightly slower than wild type cells, Fig 1C). Similarly, *lem2Δ* single mutants displayed a modest growth deficit compared to wild type, but *vps4Δlem2Δ* double mutant cells again grew much faster than *vps4Δ* cells (Fig 1C). Thus, our unbiased whole genome sequencing and targeted double mutant studies demonstrate that *cmp7* and *lem2* are bona fide *vps4* suppressors and that the spontaneous mutations phenocopy null alleles.

### *vps4∆* cells have nuclear envelope defects which are suppressed by loss of *cmp7* or *lem2*

Next, we sought to discover the cellular defects that correlated with the slow growth of *vps4Δ* cells, and to test whether those defects were also rescued by *cmp7Δ* or *lem2Δ*. In light of our prior work on mitotic and spindle pole body defects in *vps4∆* cells, we first examined nuclear envelope (NE) and spindle pole body (SPB) morphology and dynamics. Over 80% of *vps4Δ* cells from recently-dissected *vps4Δ* haploid spores displayed severe NE morphology defects. These defects were rescued by loss of either *cmp7* or *lem2* (Fig 1D-F). Moreover, approximately 30% of *vps4Δ* cells displayed clearly asymmetric SPB segregation errors and anucleate daughter cells (Fig S1A-B). In this case, loss of *cmp7* rescued these phenotypes, but loss of *lem2* did not. Indeed, even *lem2Δ* single mutant cells displayed similar SPB segregation defects (Fig S1C). Thus defects in nuclear envelope morphology and integrity were the features that correlated best with the slow growth phenotype, suggesting that these defects were primarily responsible for the *vps4Δ* slow growth phenotype.

### *vps4Δ* cells display a series of mitotic errors associated with nuclear envelope defects

Live cell imaging using NE (Ish1p-mCherry) and SPB (Cut12p-YFP) markers enabled us to monitor the development and consequences of nuclear envelope defects in mutant versus wild type cells. Abnormal NE morphologies or asymmetric and even failed karyokinesis were observed in the majority of cells (Fig 2) and only ~30% of *vps4Δ* cells displayed normal, symmetric karyokinesis (Fig 2A). An apparent proliferation or overgrowth of Ish1p-marked membranes was a particularly common defect in *vps4Δ* cells. Approximately 25% of mutant cells displayed these long-lived NE “outgrowths” that we later determined were karmellae (see below). In cells with karmellae, daughter SPBs often failed to separate normally or displayed extensive delays in separation (Fig 2B). Indeed, separation of duplicated SPBs was significantly prolonged in *vps4Δ* cells, whether or not they exhibited abnormal nuclear envelope malformations (Fig S2). Together, these observations suggest that Vps4p plays a central role in regulating NE morphology in fission yeast, particularly during SPB extrusion or insertion through the NE, and perhaps during karyokinesis.

**Figure 2.**
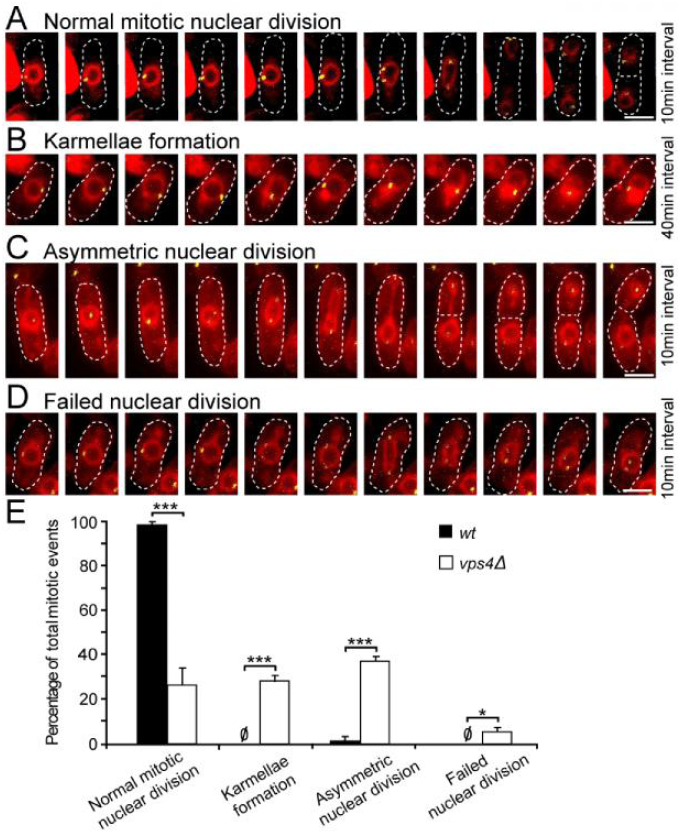
*vps4Δ* cells display a series of mitotic defects associated with failure of nuclear membrane maintenance. (A) Time lapse of normal mitotic karyokinesis in *vps4Δ* cells. In all cases, the nuclear membrane is marked by Ish1p-mCherry and the SPB is marked by Cut12p-YFP. The total length of time is 100 minutes. (B) Karmellae formation correlates with defective separation of duplicated SPB. The total length of time is 400 minutes. (C) Asymmetric nuclear division. Total length of time is 100 minutes. (D) Failed nuclear division. Total length of time is 100 minutes. (E) Quantification of NE morphology during mitosis in *wt* and *vps4Δ*. Normal mitotic nuclear division, *wt*: 99±1%, *vps4Δ*: 27±8%; Karmellae formation, *wt*: 0%, *vps4Δ*: 29±3%; Asymmetric nuclear division, Mean±SEM, *wt*: 1±1%, *vps4Δ*: 38±2%; Failed nuclear division, *wt*: 0%, *vps4Δ*: 6±2%. *wt*, n=23, 21, 18; *vps4Δ*, n=43, 51, 84. (Scale bars: 5 μm). A-D dash lines correspond with the cell wall.

### *vps4Δ* nuclei leak and their integrity is largely restored by loss of *cmp7* or *lem2*

Our observations, together with recent reports that the ESCRT pathway closes holes in the mammalian nuclear envelope (23, 24), prompted us to test the integrity of *vps4Δ* nuclei. Image analysis revealed that a large nuclear import cargo, NLS-GFP-LacZ, was enriched within nuclei by more than 10-fold in 98% of wild type cells (Fig 3A). By contrast, ~55% of *vps4Δ* cells displayed less than 10-fold nuclear enrichment (partial leaking, Fig 3B arrowheads), and ~10% of *vps4Δ* cells displayed less that 2-fold nuclear enrichment of NLS-GFP-LacZ (severe leaking, Fig 3B arrow, Fig 3C-D quantification). Remarkably, loss of *cmp7* or *lem2* rescued this abnormal nuclear integrity phenotype to a large extent, although a small minority of single and double *cmp7* or *lem2* cells still displayed partial leaking (Fig 3C-D). Live cell imaging also revealed that the extent of nuclear integrity loss correlated with NE morphology defects (Fig S3). Cells that initially displayed normal GFP reporter localization and normal NE morphology gradually accumulated cytoplasmic signal over the course of tens of minutes (Fig S3A). Cells with abnormal nuclear envelope morphology at the beginning of the experiment, by contrast, lost nuclear GFP completely over the time course (Fig S3B). Thus, cytoplasmic GFP resulted from loss of nuclear integrity rather than from defects in nuclear import.

**Figure 3.**
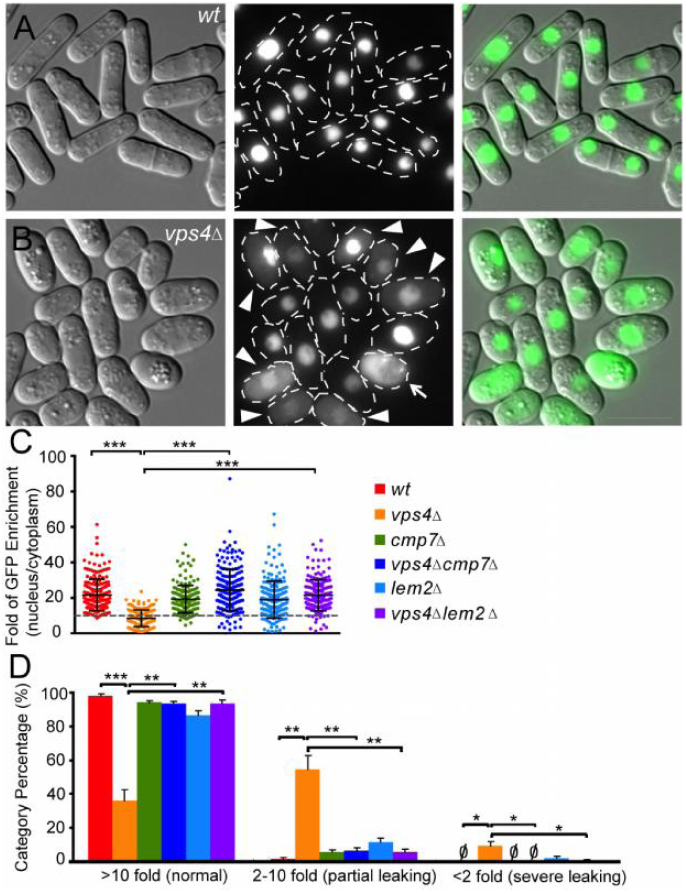
*vps4Δ* cells have leaky nuclei and nuclear integrity is restored by loss of either *cmp7* or *lem2*. (A) GFP signals in the nuclear lumen of *wt* cells expressing NLS-GFP-LacZ. (B) *vps4Δ* cells expressing NLS-GFP-LacZ with moderate (arrowheads) or severe (arrow) nuclear leaking. (Scale bar: 10 μm) A-B dash line correspond with the cell wall. (C) Nuclear enrichment of NLS-GFP-LacZ (nucleus/cytoplasm). Mean±SD, *wt*: 21.6±8.9, n=205; *vps4Δ*: 8.6±4.7, n=180; *cmp7Δ*: 19.4±7.7, n=175; *vps4Δcmp7Δ*: 24.5±11.7, n=202; *lem2Δ*: 19.1±10.4, n=198; *vps4Δlem2Δ*: 21.5±8.8, n=189. (D) Percentage of cells in different nuclear leaky phenotype categories. Mean±SEM, normal (> 10-fold GFP nuclear enrichment) *wild type*: 97.9±1.3%, *vps4Δ*: 36.2±6.5%, *cmp7Δ*: 94.1±1.1%, *vps4Δcmp7Δ*: 93.2±1.6%, *lem2Δ*: 86.6±2.3%, *vps4Δlem2Δ*: 93.4±2.0%; partial leaking (2 to 10-fold GFP nuclear enrichment) *wild type*: 1.6±0.9%, *vps4Δ*: 54.4±8.1%, *cmp7Δ*: 5.9±1.1%, *vps4Δcmp7Δ*: 6.8±1.6%, *lem2Δ*: 11.5±2.3%, *vps4Δlem2Δ*: 5.9±1.4%; severe leaking(< 2-fold GFP nuclear enrichment) *wild type*: 0%, *vps4Δ*: 9.4±2.4%, *cmp7Δ*: 0%, *vps4Δcmp7Δ*: 0%, *lem2Δ*: 2.0±1.3%, *vps4Δlem2Δ*: 0.7±0.7%; *wild type*: n=82,56,67, *vps4Δ*: n=60,59,61, *cmp7Δ*: n=62,63,50, *vps4Δcmp7Δ*: n=82,59,61, *lem2Δ*: n=68,61,69, *vps4Δlem2Δ*: n=76,50,63.

### *vps4Δ* nuclear envelopes are persistently fenestrated and have karmellae and disorganized tubular extensions

We employed electron tomography of high-pressure frozen and freeze-substituted cells to examine whether we could detect morphological defects in the nuclear envelope that could account for the loss of nuclear integrity. Serial 400nm sections were imaged and reconstructed to generate 3D volumes of >1 micron thickness. Wild type nuclei had evenly spaced inner and outer lipid bilayers with embedded nuclear pore complexes evenly distributed around the periphery (Fig 4A and Movie S1). *vps4Δ* nuclei, by contrast, displayed at least four structural abnormalities. First, large fenestrations through both the inner and outer nuclear membranes were observed (Fig 4B, bracket). Second, karmellae or concentric layers of membrane were present around certain regions of the mutant nuclei (Fig 4B, arrowheads and Fig 5B). Third, extensive whorls of disorganized tubulo-vesicular membranes that were topologically continuous with the adjacent karmellae were apparent (Fig 4B, arrows and Fig S4). Fourth, the total number of NPCs appeared to be decreased (Fig 4B, asterisk), and the NPCs that were present were localized to regions that were largely free of karmellae and tubulo-vesicular structures (Fig 4D). 3D reconstruction of these features confirmed the presence of very large gaps (>400nm) in the nuclear envelope and topologically continuous karmellae and whorls of tubular extensions (Fig 4C and Movies S2-S4). Persistent fenestrations explain the loss of nuclear integrity in *vps4Δ* cells, while karmellae and other disorganized membrane extensions may underlie the kinetic delays, asymmetries, and outright failures of SPB separation and karyokinesis.

**Figure 4.**
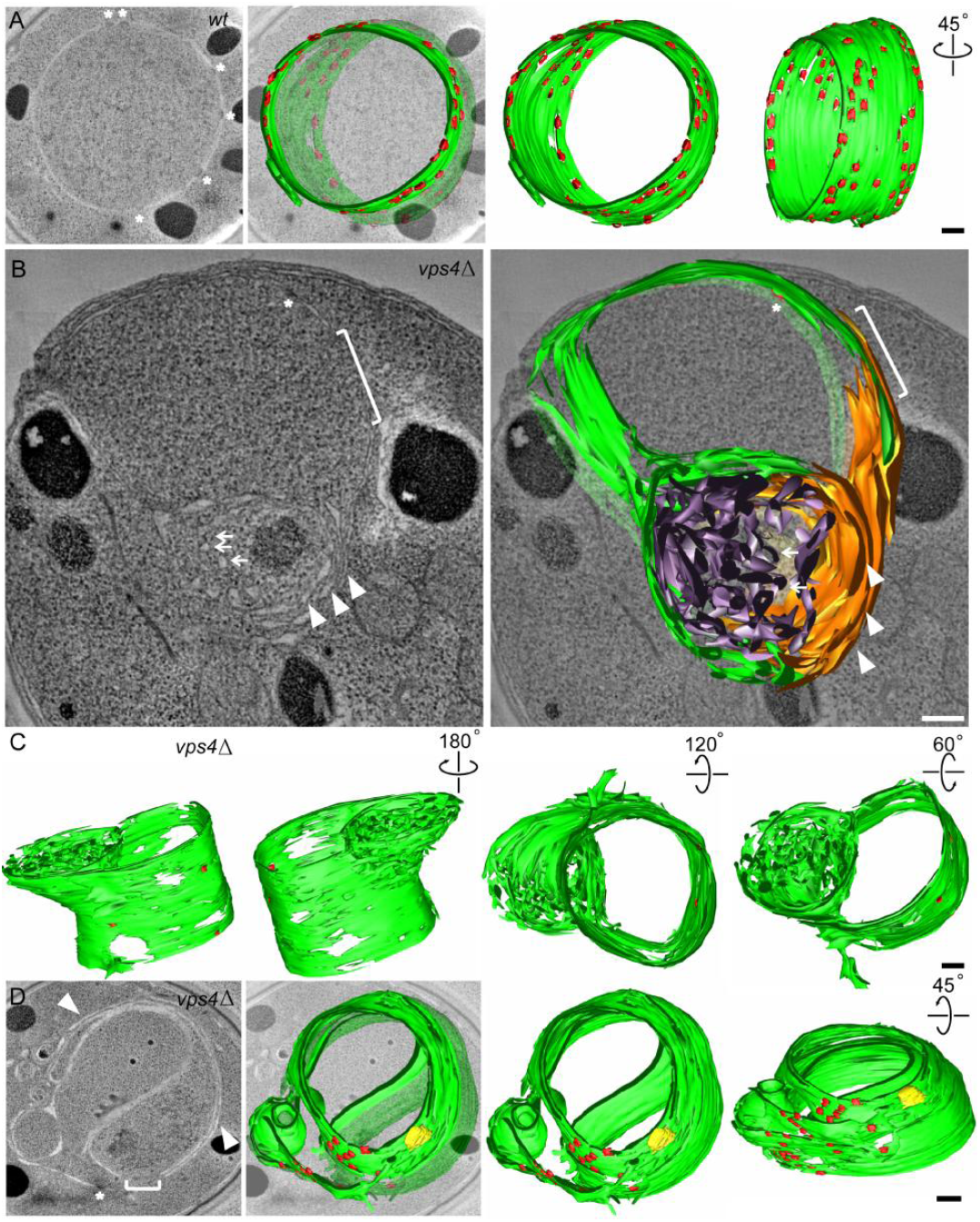
*vps4Δ* nuclear envelopes have persistent fenestrations, karmellae and disorganized tubular extensions. (A) Single tomographic slice showing the *wt* NE (left). NPCs are marked by asterisks. Segmented NE (green) with NPCs (red) reconstructed from 150 tomographic slices shown in a merged view (middle left), a front view (middle right) and a side view (right). (B) Single tomographic slice showing the *vps4Δ* NE (left). The following features and defects are highlighted: a fenestration (bracket), an NPC (asterisk), karmellae layers (white arrowheads), and a disorganized whorl of tubules (arrows). Segmented model of NE reconstructed from 100 tomographic slices is shown in a merged view (right) with karmellae in gold and a whorl of tubules in purple. (C) A segmented model of *vps4Δ* NE reconstructed from 200 tomographic slices is shown from the front (left), back (middle left), top (middle right) and bottom (right). (D) Single *vps4Δ* tomographic slice showing NPCs are absent from karmellae region (left). The following features and defects are highlighted: a fenestration (bracket), an NPC (asterisk), and karmellae layers (white arrowheads). Segmented NE (green) with NPCs (red) and SPB (yellow) reconstructed from 150 tomographic slices shown in a merged view (middle left), a front view (middle right) and a side view (right). (Scale bars: 200 nm).

**Figure 5.**
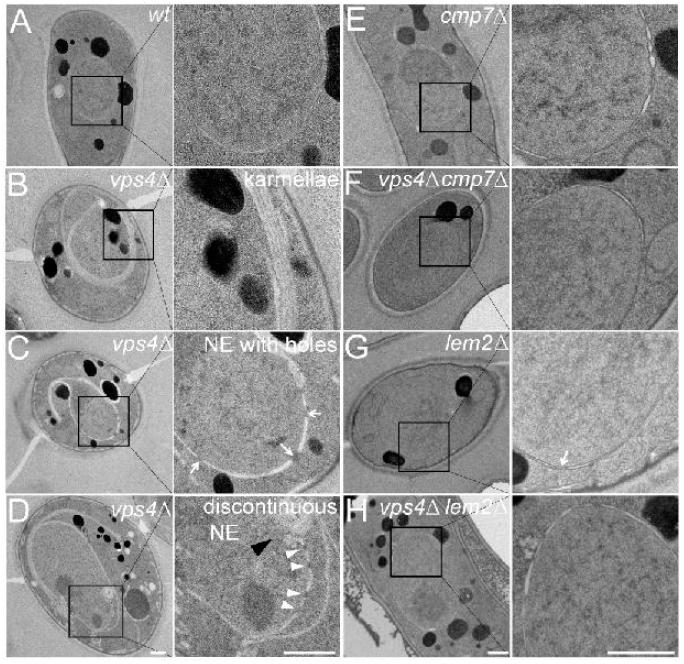
*vps4Δ* nuclear envelopes have persistent fenestrations, karmellae and disorganized extensions, and these defects are suppressed by loss of *cmp7* or *lem2*. (A) *wt* cells with normal NE. (B-D) *vps4Δ* cells with a variety of NE phenotypes, including karmellae (B), multiple holes (arrows) (C), and discontinuities (black arrow head) that can contain apparently intra-nuclear NPCs (arrowheads, D). (E and G) *cmp7Δ* and *lem2Δ* single mutants versus (F and H) *vps4Δcmp7Δ* and *vps4Δlem2Δ* double mutants. (Scale bars: 500 nm).

### *vps4Δ* nuclear envelope defects are suppressed by loss of *cmp7* or *lem2*

Thin section electron microscopy of single and double mutant cells demonstrated that karmellae formation in *vps4Δ* cells was completely suppressed by loss of *cmp7* or *lem2* (Fig 5). Remarkably, double mutant cells displayed wild-type like nuclei, although a few examples of probable nuclear fenestrations were observed in *lem2Δ* single mutant cells (Fig 5G). These observations indicate that in the absence of Vps4p, karmellae formation depends on both Lem2p and Cmp7p. Similarly, overexpressing Lem2p in *S. pombe*, or compromising nuclear import of Heh2p, an orthologue of Lem2p in *S. cerevisiae*, induces the formation of similar abnormalities (31, 32). Together, these results suggest that in the absence of *vps4*, unregulated Lem2p activity drives formation of toxic nuclear envelope malformations via a pathway that also requires Cmp7p (Fig S5).

### LEM2 recruits CHMP7 to the reforming nuclear envelope during anaphase in mammalian cells

LEM2 and its homologs are two-pass membrane proteins that reside in the inner nuclear membrane (32, 33). A LEM2 homolog in budding yeast, Heh2, has been previously implicated in ESCRT-dependent surveillance of defective NPCs (21). These studies and our observations suggested that LEM2-like proteins in both budding and fission yeast may be site-specific nuclear membrane adaptors that recruit and/or activate Cmp7p/CHMP7. While a role for CHMP7 in nuclear envelope closure has been reported in mammalian cells (24), its localization during the process of nuclear assembly had not been determined. To confirm that CHMP7 is indeed present at this site and to track the spatial relationship between LEM2 and CHMP7 in human cells, we assessed the dynamics of LEM2-mCherry and GFP-CHMP7 localization by live imaging of HeLa cells. Like other nuclear envelope proteins that reside in the endoplasmic reticulum during mitosis (34), LEM2-mCherry was found in a reticular network as cells emerged from metaphase. Early in anaphase, LEM2-mCherry rapidly accumulated around condensed chromatin disks - first at distal ends, then concentrating at the central or “core” region of the anaphase disk and then broadening to a distributed nuclear rim pattern. The latter pattern has been reported previously (35). CHMP7 recruitment to the chromatin surface coincided closely in time with the appearance of LEM2, resulting initially in their robust concentration at the same regions of the chromatin disks (Fig 6A, accompanying Movie S5; Fig S6A). The spatiotemporal specificity of the CHMP7 and LEM2 recruitment during early anaphase contrasts with their lack of colocalization after cleavage furrow ingression (Fig 6A, Movie S5, Fig S6A) and their even more distinct spatial distributions during interphase, when GFP-CHMP7 was no longer present at the chromatin surface and LEM2-mCherry decorated the nuclear rim as expected (Fig S6B).

**Figure 6.**
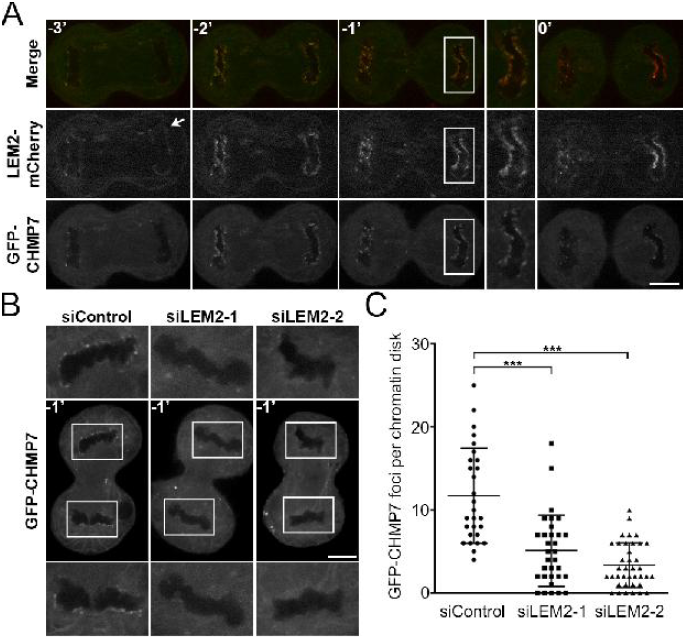
Live imaging of LEM2-dependent recruitment of CHMP7 to reforming nuclei during mammalian anaphase. (A) Montage of a representative cell expressing LEM2-mCherry and GFP-CHMP7 progressing through anaphase before complete furrow ingression (designated as t=0’). LEM2-mCherry makes initial contacts with chromatin (white arrow) and GFP-CHMP7 localizes to sites of LEM2-mCherry enrichment (t=−2’, −1’). (B) Illustrative cells, at t=−1’, treated with siRNA as indicated and expressing GFP-CHMP7. (C) Quantification of GFP-CHMP7 peaks at chromatin disks of cells treated with siRNA (Mean±SD; siControl: 12±6%, n=28; siLEM2-1: 5±4%, n=32; siLEM2-2: 3±3%, n=38). (Scale bars: 10 μm).

We next tested whether LEM2 is required for CHMP7 recruitment to reforming nuclei. Following depletion of endogenous LEM2 with two independent siRNA oligos (Fig S7A-C), GFP-CHMP7 recruitment to anaphase chromatin was notably attenuated (Fig 6B-C; Fig S8). These results were also recapitulated in a cell line expressing H2B-mCherry, confirming that CHMP7 foci are in close apposition to the chromatin surface and that this localization is impaired following LEM2 depletion (Movies S6-S7). To further evaluate the role of LEM2 in recruiting the ESCRT pathway to the reforming nuclear envelope, we also tracked the localization of IST1/CHMP8 and CHMP2A, two ESCRT-III proteins that were previously shown to be recruited to the surface of condensed chromatin disks to promote coordinated NE closure and disassembly of spindle microtubules (24) (also see Movie S8). siRNA depletion of CHMP7 abrogated robust IST1/CHMP8 and CHMP2A recruitment to chromatin as assayed by detection of endogenous protein by fixed cell immunofluorescence (Fig 7B, 7D; Fig S7D, S7G; Table S2 and S3), consistent with the previously reported role for CHMP7 in the recruitment of CHMP4B to assembling nuclei (24). Importantly, siRNA-mediated depletion of LEM2 also strongly attenuated robust IST1/CHMP8 and CHMP2A recruitment (Fig 7A-D; Table S3 and S4). By contrast, depletion of the abundant inner nuclear membrane (INM) protein SUN1 did not alter IST1/CHMP8 localization (Fig S7E-G), suggesting a specific requirement for LEM2 in this pathway. To clarify the epistatic relationship between CHMP7 and LEM2, we also depleted CHMP7 and measured LEM2 recruitment to chromatin disks during anaphase. Loss of CHMP7 had no effect on level of LEM2 recruitment (Fig 7E and F, and Table S4).

**Figure 7.**
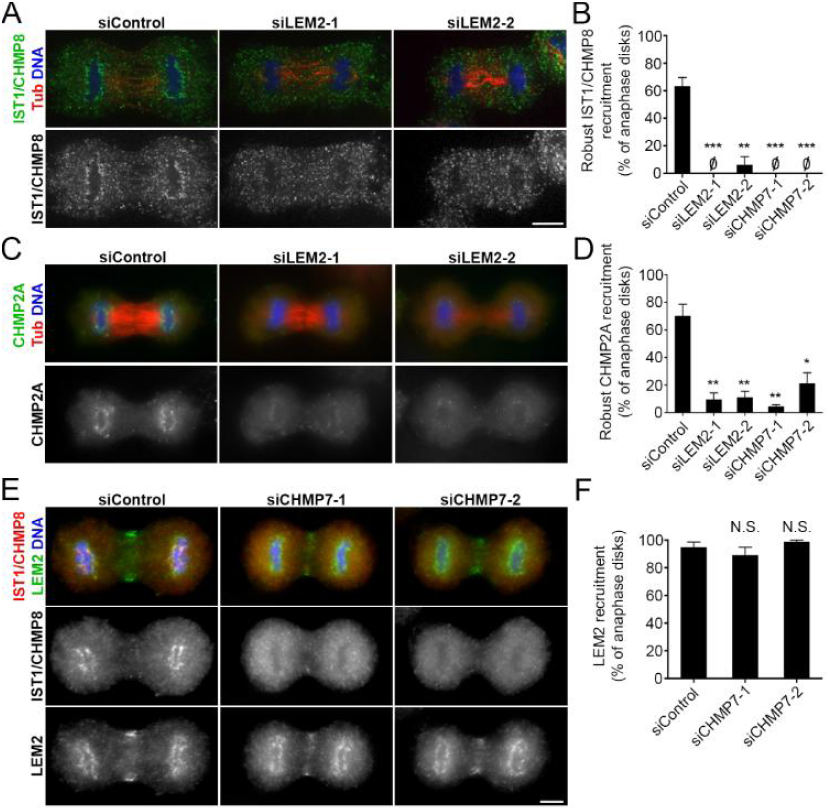
Recruitment of CHMP2A and IST1/CHMP8 during mammalian nuclear reformation depends on LEM2, whose targeting is independent of CHMP7. (A) Confocal images illustrating IST1/CHMP8 localization in anaphase B cells after siControl, siLEM2-1, or siLEM2-2 treatment. (B) Quantification of IST1/CHMP8 recruitment to chromatin disks during anaphase B. IST1/CHMP8 recruitment was scored as robust, weak, or no chromatin-associated foci. The robust category is graphed and statistical analysis performed, comparing the siControl dataset to each depletion condition dataset (siControl: 63±6%, n=18, 58, 24; siLEM2-1: 0±0%, n=40, 44, 12; siLEM2-2: 6±6%, n=34, 22, 6; siCHMP7-1: 0±0%, n=42, 20, 12; siCHMP7-2: 0±0%, n=22, 20, 2). (C) Widefield images illustrating CHMP2A localization in anaphase B cells after siControl, siLEM2-1, or siLEM2-2 treatment. (D) CHMP2A recruitment to chromatin disks at anaphase B, assessed as in B, above (siControl: 70±9%, n=48, 48, 52; siLEM2-1: 9±5%, n=108, 86, 38; siLEM2-2: 11±5%, n=98, 52, 42; siCHMP7-1: 4±1%, n=78, 102, 47; siCHMP7-2: 21±8%, n=112, 58, 56). Analysis of parallel samples confirmed that LEM2 and CHMP7 depletion profoundly disrupted IST1/CHMP8 recruitment as before and is shown in supplemental Fig S7G. (E) Widefield images of cells co-stained for IST1/CHMP8 and LEM2 illustrates the differential sensitivity of their localization at anaphase chromatin disks after siCHMP7-1 or siCHMP7-2 treatment. Signal detected at the midzone with LEM2 antibody is likely non-specific, see supplemental Fig S7A. (F) Quantification of LEM2 recruitment to chromatin disks at anaphase B. Images were used for blind scoring the presence of chromatin-associated LEM2 (siControl: 95±4%, n=28, 35, 39; siCHMP7-1: 90±6%, n=18, 72, 45; siCHMP7-2: 99±1%, n=35, 56, 7). Analysis of parallel samples confirmed that CHMP7 depletion profoundly disrupted IST1/CHMP8 recruitment as before and is shown in supplemental Fig S7D. (Scale bars: 10 µm). All graphs plot Mean±SEM.

Finally, to determine whether LEM2 can bind directly to CHMP7, we purified full-length human CHMP7 as well as the soluble N-terminal versus C-terminal domains of human LEM2 to homogeneity. After immobilization on Ni-NTA beads via a fused His-SUMO tag, the C-terminal domain—but not the N-terminal domain—bound full-length CHMP7 (Fig S9). Thus, our imaging, siRNA depletion and biochemical data all support the idea that LEM2 binds CHMP7 directly and serves as an adaptor that recruits CHMP7 and other downstream ESCRT-III proteins, including IST1/CHMP8 and CHMP2A, to the reforming nuclear envelope during anaphase.

## Discussion

Pioneering work on the ESCRT pathway in budding yeast led to our understanding of its roles in multi-vesicular body biogenesis (18), while work in human cells is leading to a broader view of ESCRT roles at a variety of different target membranes (20). Our results indicate that the role of the ESCRT pathway in closing holes in the nuclear envelope is evolutionarily ancient ((16) and this study). The different nuclear ESCRT functions (NPC surveillance, nuclear envelope reformation, and SPB insertion/removal) can now be unified by the hypothesis that all of these processes generate fenestrations in the nuclear envelope that are closed by the action of the ESCRT machinery ((21, 23-26, 36) and this study). Conserved activities include roles for nucleus-specific adaptors of the LEM domain family and the ESCRT-II/ESCRT-III hybrid protein, CHMP7/Cmp7p (21, 29) and this study). While our study was under review, corroborative evidence for these roles for CHMP7 in human cells (37) and LEM domain family proteins in budding yeast were also published by others (38). This conserved pathway may likewise underlie the requirement for Src1, a LEM-domain protein in *A. nidulans*, in the formation of stable nuclei (39). This body of work suggests that LEM2 plays a specific, initiating role in coordinating membrane remodeling events, particularly during nuclear assembly, in addition to the other roles it plays as a nuclear envelope resident during interphase (31, 35, 40-45).

Two LEM family members, Heh1p and Heh2p, are involved in ESCRT recruitment in *S. cerevisiae* have, so multiple LEM family proteins may also be involved in recruiting ESCRT-III activities in mammalian cells. In this regard, we found that knockdown of LEM2 in HeLa cells led to a less severe IST1/CHMP8 recruitment phenotype than knockdown of CHMP7 (Table S2 and S3). It will therefore be of great interest to determine whether additional LEM-domain family members, which are present at the nascent nuclear envelope (46, 47), also serve as ESCRT recruitment factors in human cells. Like other ESCRT-III proteins, CHMP7 can also interact with membranes, and this activity contributes to its targeting (37), so further biochemical studies will be required to elucidate how the dynamic interplay between LEM2, CHMP7 and lipids regulates the recruitment and activity of the ESCRT pathway at the nascent nuclear envelope. It will additionally be informative to determine the mechanistic basis of *vps4Δ* phenotypes in fission yeast and the underlying toxicity of unrestricted Lem2p-Cmp7p activity. We speculate that in the absence of Vps4p, Lem2p-Cmp7p complexes stabilize nuclear membrane fenestrations aberrantly and/or promote nuclear envelope remodeling events that—when unregulated by Vps4p—result in karmellae, large gaps, and other malformations that ultimately compromise cell division (Fig S5). Similarly, over-expression of Lem2p in fission yeast induces nuclear envelope malformations (31); and impairment of Heh1/2p nuclear import in budding yeast induces karmellae formation (32). We predict that both of these phenotypes depend on CHMP7/Cmp7p proteins and other downstream ESCRT pathway activities. Studies of these and related activities in the future will benefit from the facile *S. pombe* genetic system for investigating how the ESCRT pathway senses and seals breaches of the envelope—a nuclear membrane integrity pathway that is conserved from yeast to human.

## Materials and Methods

### Yeast strains and growth media

*S. pombe*, diploid strain SP286 (*h+/h+, leu1-32/leu1-32, ura4-D18/ura4-D18, ade6-M210/ade6-M216*) was used for all haploid constructions. All other strains used in this study are listed in Appendix Table S1 and S5. Cells were routinely grown in YE5 rich media (yeast extract 0.5% w/v, glucose 3.0% w/v supplemented with 225 mg/L histidine, leucine, uracil and lysine hydrochloride and 250 mg/L adenine) or Edinburgh Minimal Media (EMM, Sunrise Sciences Products) with supplements described above (EMM5). Sporulation of diploids was induced by culturing cells in EMM5 without glutamate (EMMG, Sunrise Sciences Products) and supplemented as above but without uracil. Dominant drug selection was performed with YE5 supplemented with G418 disulfate (KSE Scientific) at 200 mg/L, hygromycin B, *Streptomyces, sp.* (Calbiochem) at 100 mg/L and ClonNAT at 100 mg/L. To prevent caramelization, YE5 was routinely prepared by leaving out glucose during autoclaving and adding it before inoculation.

### Yeast transformation

Log phase yeast cells were incubated in 0.1M Lithium Acetate (pH 4.9) for 2 hours, at a concentration of 5×10^8^ cells/ml. 100 μl of this cell suspension was then mixed with 0.5-1 μg DNA and 290 μl 50%(wt/vol) PEG8000 and incubated overnight. Cells were recovered in 0.5X YE5 medium overnight before plating. All steps were conducted at 32°C.

### Sporulation, random spore analysis and tetrad dissection

Sporulation of *S. pombe* diploids for tetrad dissection or random spore analysis was induced by first transforming pON177 (*h-* mating plasmid with ura4 selectable marker, a gift from Megan King, Yale School of Medicine) into the parental strain. Transformants were selected on EMM5 without uracil (EMM5-uracil) for three days at 32°C followed by induction in 0.5 ml EMMG without uracil (EMMG-uracil) for 36 to 48 hours at 25°C. Sporulation was confirmed by microscopy. Ascus walls of tetrads were digested with 2% (v/v) β-glucuronidase (Sigma-Aldrich) overnight at 25°C. This overnight treatment also eliminated non-sporulated diploids. Enrichment of spores was verified by microscopy. Five to 10 fold serial dilutions were made and spores were plated on YE5 supplemented with specific antibiotics.

Tetrad dissection: 10-fold serial dilutions of β-glucuronidase-treated tetrads were placed onto YE5 plates as a narrow strip and digestion was monitored at room temperature. Tetrads were picked and microscopically dissected along pre-designated lines of the same YE5 plate. Spores were allowed to germinate and grow at 32°C.

### Knockout cassettes and plasmids

*vps4* deletion cassette: the *vps4Δ::natMX6* template (16) was amplified to create an amplicon covering 756 bp upstream of the ATG and 499 bp downstream of the stop codon. This 2439 bp amplicon was transformed into SP286 and plated on YE5 + ClonNAT to select for heterozygous *vps4Δ::natMX6/+* diploids.

*cmp7* deletion cassette: a fragment including 420 bp upstream of the ATG and 377 bp downstream of the stop codon was amplified and cloned into BamHI/BglII and EcoRV/SpeI of pAG32 (a pFA6 derived plasmid containing *hphMX4,* which confers resistance to hygromycin B, was a gift from David Stillman, Department of Pathology, University of Utah). The *cmp7Δ::hphMX4* fragment was amplified and transformed into heterozygous *vps4Δ::natMX6/+* diploid intermediate strain and selected on YE5 supplemented with both ClonNAT and hygromycin B.

*lem2* deletion cassette: a 3.1 kb *lem2* genomic fragment spanning 600 bp upstream of the ATG to 500 bp downstream of the stop codon, was subcloned into pGEM vector. The ORF region was replaced with a *hphMX4* hygromycin resistance cassette to make the final knock-out construct (pMGF130). The *lem2Δ::hphMX4* cassette was amplified and transformed into heterozygous *vps4Δ::natMX6/+* diploids and selected on YE5 supplemented with both ClonNAT and hygromycin B.

*cut12-YFP* cassette: a 3.5 kb synthetic DNA fragment was created that spanned 550 bp C-terminal of *cut12* fused with *YFP* followed by *kanMX6* and a 500 bp fragment downstream of the *cut12* stop codon. The fragment was transformed into heterozygous diploid strains and the resulting diploids were selected on YE5 supplemented with G418 disulfate.

*ish1-mCherry* cassettes: a 1kb *ish1* genomic DNA fragment corresponding to 500 bp upstream and 500 bp downstream of the *ish1* stop codon was subcloned into the pGEM vector. A flexible linker (GGTGGSGGT) and *mCherry* fusion cassettes were assembled, followed by the *yADH1* terminator and MX4/6 drug resistance or auxotrophic markers. These cassettes were integrated at the native *ish1* locus to make the final fusion constructs: *ish1-mCherry::natMX6* (pMGF170), *ish1-mCherry*::*kanMX6* (pMGF169), *ish1-mCherry*::*hphMX4* (pMGF157) and *ish1-mCherry*::*ura4(+)* (pMGF172).

*pDual-SV40NLS-GFP-LacZ* construct: a *SV40NLS-GFP-LacZ* fragment was amplified from pREP3X (provided by Dr. Shelley Sazer, Baylor College of Medicine) and subcloned downstream of the *Pnmt1* promoter in the *pDual* vector. The final construct (pMGF173) was integrated at the *leu1* locus.

*Pcmv(Δ5)*-*GFP-CHMP7* and *pCMV(Δ5)*-*LEMD2-mCherry*: *CHMP7* and *LEMD2* cDNA were amplified and sublconed downstream of Pcmv(*Δ5)* (48). GFP+linker (see as above) was inserted before *CHMP7* (pMGF182) and linker+mCherry was inserted after *LEMD2* (pMGF196) to make the fusions.

### Isolation of *vps4Δ* and suppressors

Isolation and handling of *vps4Δ* haploids: individual petite colonies from random spore analysis plates of YE5+ClonNAT were selected, re-suspended in 200 μL YE5 media, plated onto two YE5+ClonNAT plates and incubated at 32°C for three days. Isolates that grew across one plate without apparent suppression were frozen with glycerol without further culturing by scraping and re-suspension in YE5 with 15%(wt/vol) glycerol. Cells from the other plate were scraped and genomic DNA was immediately extracted for Illumina sequencing (see below).

Isolation of *vps4Δ* suppressors: *vps4Δ* isolates were re-streaked from glycerol stocks and cultured on YE5 at 32°C. Large colonies (apparent suppressors) were picked, resuspended in 200 μL YE5 media and plated onto two YE5 plates. After two days, cells were harvested as described above for glycerol stocks and genomic DNA extraction.

### Yeast genomic DNA extraction, Illumina sequencing and analysis

Genomic DNA extraction: frozen pellets of *wt*, *vps4Δ* or suppressor cells (200 μl) were thawed on ice. 250 μL resuspension buffer (20 mM Tris-HCl, pH 8.0, 100 mM EDTA, 0.5 M β-mercaptoethanol) and 50 μL lyticase (50 units)) were then added to remove the cell wall. Genomic DNA was extracted using phenol/chloroform/isoamyl alcohol, precipitated with ethanol and treated with RNase, followed by DNeasy Blood Tissue purification according to the manufacturer’s protocol (Qiagen cat. 69504).

Illumina sequencing: libraries were constructed using the Illumina TruSeq DNA Sample Preparation Kit (cat. no. FC-121-2001, FC-121-2002). Briefly, genomic DNA was sheared in a volume of 52.5 μl using a Covaris S2 Focused-ultra-sonicator with the following settings: Intensity 5.0; Duty cycle 10%; Cycles per Burst 200; Treatment time 120 seconds. Sheared DNA was converted to blunt-ended fragments and size selected to an average length of 275 bp using AMPure XP (Beckman Coulter cat. no. A63882). Following adenylation of the DNA, adapters containing a T-base overhang were ligated to the A-tailed fragments. Adapter ligated fragments were enriched by PCR (8 cycles) and purified with AMPure XP. The amplified libraries were qualified on an Agilent Technologies 2200 TapeStation using a D1K ScreenTape assay (cat. no. 5067-5363) and quantitative PCR was performed using the Kapa Biosystems Kapa Library Quant Kit (cat. no. KK4824) to define the molarity of adapter-modified molecules. Molarities of all libraries were subsequently adjusted to 10 nM and equal volumes were pooled in preparation for Illumina sequencing.

Sequencing data analysis: raw reads were aligned to the *S. pombe* genome, obtained from Ensembl Fungi, using NovoCraft Novoalign allowing for no repeats (-r None) and base calibration (-k). Alignments were converted to Bam formatted files using samtools (http://samtools.sourceforge.net). Sequence pileups were generated with samtools pileup, and variants called using the bcftools utility (options -c -g -v). Variants were filtered using the included varFilter Perl script included with samtools and written out as a vcf file. To distinguish unique variants in each strain from common variants, sample vcf files were intersected with one another using the Perl script intersect_SNPs (https://github.com/tjparnell). Variants were annotated with the Perl script locate_SNPs (https://github.com/tjparnell) using a GFF3 gene annotation file obtained from Ensembl. From the resulting table, variants were further filtered by the fraction of reads supporting the alternate allele, the presence of codon changes, and visual inspection in a genome browser. The summary statistics are reported in Table S1.

### Fluorescence microscopy

All yeast strains were cultured in either YE5 or in EMM5 if the desired protein was induced by the *nmt1* promoter. Cells were imaged after reaching log phase. Hoechst staining was conducted at 1μg/ml in water for 15 minutes. Images were collected on a Zeiss Axio Observer Z1 microscope using a 100X oil-immersion objective.

HeLa cells were fixed in −20°C methanol for 10 minutes. The primary antibodies used for immuno-detection were rabbit α-IST1/CHMP8 (49), α-LEM2 (HPA017340; Sigma), rat α-tubulin (YL1/2; Accurate Chemical & Scientific), rabbit α-CHMP2A (UT 634; Covance) and mouse α-IST1/CHMP8 (UT 697; Covance). Full length CHMP2A and IST1/CHMP8 protein (50) were used as antigens to produce custom antibodies by Covance Immunology Services. The anti IST1/CHMP8 antibody (UT 697) was affinity purified (51) before use. Following incubation with fluorescently labeled secondary antibodies (Thermo Fisher), coverslips were mounted using DAPI ProLong Gold (Thermo Fisher) and imaged. For the purpose of illustration, images of anaphase B cells were acquired by spinning disk confocal microscope and adjusted so that cytoplasmic IST1/CHMP8 intensity was comparable between samples. Images acquired by widefield microscopy at 100X were used to score the IST1/CHMP8 and CHMP2A phenotypes following minimal adjustment that was applied uniformly. For each graph, IST1/CHMP8 and CHMP2A localization to anaphase B chromatin masses were assessed in three independent experiments. Each chromatin disc (two per cell) was scored as having either robust, weak or no recruitment for IST1/CHMP8 or CHMP2A. Robust recruitment was characterized by distinctive foci organized at chromatin masses while weak recruitment was characterized by less intense, often fewer and less organized foci at the chromatin surface. Images of anaphase B cells from all treatments and experiments were randomized and quantified blindly by three scorers. The majority score was used in cases where the three scores differed. In an analogous set of experiments, anaphase B cells from all treatments were scored blindly for the absence or presence of chromatin-associated LEM2. As a positive control, anaphase B cells were scored, in parallel, as having either robust, weak or no IST1/CHMP8 recruitment.

### Time-lapse light microscopy analysis

Wild type and *vps4Δ* cells were grown in YE5 at 32°C for 8 hours and loaded into the CellASIC ONIX Microfluidic system (Cat. EVE262, EMD Millipore), which immobilizes the cells in a single focal plane, maintains a constant temperature (32°C) and pumps fresh media over the cells. Images from multiple positions per chamber were captured every 10 minutes for 16 hours. A lens heater was used to maintain constant temperature inside the chamber. Images were collected with an Andor Clara CCD camera attached to a Nikon Ti microscope using a 60x oil Nikon Apo Lambda S NA 1.4 lens. The samples were illuminated with a Lumencor Sola LED at 20% intensity, which was further reduced by the insertion of an ND8 filter. Exposure times ranged between 1 to 3 seconds for both YFP and mCherry channels. Five Z plane images separated by 1 micron were collected. Maximum intensity projection images were created to follow the Cut12p-YFP signals within a given cell. For Ish1p-mCherry signals, only the Z plane that bisected the nucleus was chosen for further image analysis.

For time-lapse co-localization experiments in HeLa cells, cells were plated on fibronectin-coated Mat-Tek dishes and incubated overnight. Cells were then transiently co-transfected with pCMV(Δ5)-GFP-CHMP7 and pCMV(Δ5)-LEM2-mCherry using Lipofectamine 3000 (Thermo Fisher) to co-express GFP-CHMP7 and LEM2-mCherry under attenuated CMV promoters (48). For siRNA depletion and GFP-CHMP7 expression experiments, HeLa cells, either parental or stably expressing H2B-mCherry, were plated on fibronectin-coated Mat-Tek dishes in the presence of siRNA (siControl, siLEM2-1, or siLEM2-2, as described below). After 24 hours, pCMV(Δ5)-GFP-CHMP7 was delivered by transient transfection with Lipofectamine 3000. In all live imaging experiments, cells were transiently transfected for 24 hours before being arrested at G1/S and released, as described below. Twelve hours after release, cells were live-imaged by spinning disk confocal microscopy.

### Quantification of nuclear enrichment of NLS-GFP-LacZ

Fluorescence microscopy of fission yeast was performed as described above using 0.2s exposures for 5 Z sections separated by 0.29μm steps. Integrated pixel intensities for GFP were measured within a 100×100 pixel square box at both the center of nucleus and cytoplasm near the pole of the cell. The average background pixel intensity was also measure from cell-free regions of the image, and this value was subtracted from both the nuclear sum and the cytoplasmic sum. The fold nuclear GFP enrichment was calculated as the ratio of nuclear/cytoplasmic integrated intensity. A minimum of 150 cells were quantified for each genotype.

### Quantification of CHMP7 recruitment during anaphase

Live cell fluorescence microscopy was performed as described above. Images of cells from 1 minute before cleavage furrow ingression were selected for scoring. The “Find Maxima” function with variable noise tolerance values, as implemented in Image J, was used to identify CHMP7 foci around the contour of chromatin disks in each cell. The absolute number of those foci were recorded and scored blindly between siControl and siLEM2.

### Electron microscopy

Yeast strains were grown to log phase and harvested using gentle vacuum filtration onto filter paper. The cell pellet was scraped from the filter, mixed with cryo-protectant (20%(wt/vol) BSA in PBS), transferred to the well of a 100 µm specimen carrier (type A), and covered with the flat side of a type B specimen carrier (52). The loaded specimens were immediately frozen with a high-pressure freezer (EM-HPM100; Leica Microsystems, Vienna). Frozen cell were transferred for freeze-substitution (FS) to a pre-cooled Leica AFS unit (Leica Microsystems) and processed in the following substitution solution: 97% anhydrous acetone (EMS, RT10016) and 3% of water, 2% osmium tetroxide and 0.1% uranyl acetate. Substitution started at –90ºC for 72 hours followed by a gradual increase in temperature (5ºC/hr) to –25ºC over 13 hours, held at - 25ºC for 12 hours, then warmed (10ºC/hr) to 0ºC over 2.5 hours). The samples were removed from the AFS unit, washed 6 times with pure acetone, and gradually infiltrated and embedded in Epon12-araldite resin as follows: 50%(vol/vol) epon12-araldite/acetone overnight, 75%(vol/vol) epon12-araldite/acetone for 24 hours, 100% epon12-araldite for 8 hours and polymerized at 60°C for 48 hours. Ultrathin sections (70 nm) were cut using a diamond knife (Diatome, Switzerland), using a UC6 microtome (Leica Microsystems), transferred to Formvar- and carbon-coated mesh copper grids (Electron Microscopy Sciences, FCF-200-Cu) and post-stained with 3%(wt/vol) uranyl acetate and Reynold’s lead citrate. The sections were viewed with a JEM-1400 Plus transmission electron microscope (JEOL, Ltd, Japan) at 120 kV and images collected on a Gatan Ultrascan CCD (Gatan Inc., Pleasanton CA).

### Electron tomography

Blocks of embedded *wt* and *vps4Δ S. pombe* cells were trimmed to ~100 × 200 μm faces. Serial semi-thick (400 nm) sections were cut with a UC6 ultramicrotome (Leica Microsystems) using a diamond knife (Diatome Ltd. Biel, Switzerland). Ribbons of 10-20 sections were placed on Formvar-coated, copper-rhodium 1 mm slot grids (Electron Microscopy Sciences, Fort Washington PA) and stained with 3%(wt/vol) uranyl acetate and Reynold’s lead citrate. Colloidal gold particles (10 nm) were placed on both surfaces of the sections to serve as fiducial markers for subsequent tomographic image alignment and the grids carbon coated to enhance stability in the electron beam.

Grids were placed in a dual-axis tomography holder (Model 2040; EA Fischione Instruments, Inc., Export, PA.) and imaged with a Tecnai TF30-ST transmission electron microscope (FEI Company, Hillsboro, OR) equipped with a field emission gun and operating at 300 kV. Well-preserved cells displaying structures of interest were identified and tracked over 6-10 serial sections. The volume of the cell present in each section was then imaged as a dual-axis tilt series; for each set, the grid was tilted +/-64° and images recorded at 1° intervals. The grid was then rotated 90° and a similar series recorded about the orthogonal axis. Tilt-series datasets were acquired automatically using the SerialEM software package (53) and images recorded with a XP1000 CCD camera (Gatan Inc., Pleasanton CA). Tomographic data was calculated and serial tomograms were joined together using the IMOD software package (54-56). Tomograms were analyzed, segmented and modeled using IMOD. The nuclear envelope was traced with closed contours in each tomogram. Modeled contours were smoothed and 3D surfaces generated with tools in the IMOD software package. The “Z inc” value was set to 3 for all nuclear envelope objects to further smooth their surface. The number of serial semi-thick (400nm) sections used for model segmentation and display were: 3 for *wt* (Movie S1); 2 for *vps4Δ* (Movie S2); 4 for *vps4Δ* (Movie S3) 3 for the second *vps4Δ* (Movie S4).

### siRNA-mediated depletion and cell cycle synchronization

HeLa cells were plated on fibronectin-coated coverslips in the presence of 10 nM siRNA oligo, delivered by Lipofectamine RNAiMAX Transfection Reagent (Thermo Fisher). Specific sequences used were: siControl [siScr-1; (57)], siLEM2-1 [antisense sequence targeting nucleotides 78-98: UUGCGGUAGACAUCCCGGGdTdT; (45)], siLEM2-2 [antisense sequence targeting nucleotides 1297-1317: UACAUAUGGAUAGCGCUCCdTdT; (45)], siCHMP7-1 [CHMP7 650; sense sequence: GGGAGAAGAUUGUGAAGUUdTdT; (22)], and siCHMP7-2 [CHMP7 613; sense sequence: CAGAAGGAGAAGAGGGUCAdTdT; (22)], siSUN1A [SUN1; sense sequence targeting nucleotides 2321-2345 of SUN1, transcript variant 1 (NM_001130965.2): CCAUCCUGAGUAUACCUGUCUGUAUdTdT; (58)], and siSUN1B [SUN1; sense sequence targeting nucleotides 865-883 of NM_001130965.2: UUACCAGGUGCCUUCGAAAdTdT; (59)]. Culture media containing 2 mM thymidine was then added for 24 hours to arrest cells at G1/S. Cells were then rinsed thoroughly with PBS followed by the addition of culture media. Twelve hours after release, cells were imaged live or fixed for microscopy. To verify efficacy and specificity of siRNA treatments, HeLa cells were plated in 6-well dishes and subjected to similar conditions as above. Cell lysates were then harvested and analyzed with immunoblots. Following incubation with primary antibodies (α-LEM2 (HPA017340; Sigma), α-CHMP7 (HPA036119; Sigma), α-tubulin (YL1/2), α-SUN1 (EPR6554; Abcam)), reactivity was detected using HRP-coupled secondary antibodies (Thermo Fisher) and chemiluminescence.

### Protein Expression and Purification

We individually expressed full length human CHMP7 (Uniprot ID Q8WUX9) and the terminal domains of LEM2, encoded by *LEMD2* (NTD 1-208, CTD 395-503, Uniprot ID: Q8NC56) with a N-terminal His_6_-Sumo affinity tag using a pCA528 vector (WISP08-174, DNASU Plasmid Repository) in BL21-(DE3)-RIPL *E. coli*. The cells were grown in auto-induction media ZYP-50529 (1.5 L cultures each). Cells were grown at 37ºC to OD 0.8 with vigorous shaking in baffled flasks, moved to 19ºC and grown for an additional 19 hours. Cells were then harvested by centrifugation and bacterial pellets were snap-frozen in liquid nitrogen. Subsequent purification steps were performed at 4ºC. Cells were thawed and lysed for 30 min with lysozyme in 2.5 times the pellet volume of lysis buffer followed by sonication. The supernatant was clarified by centrifugation (40,000 *g*, 45 mins).

*CHMP7 purification:* 20 to 30 g of bacterial pellet were lysed (50 mM Tris, pH 8.0, 1 M NaCl, 20 mM Imidazole, 10% (wt/vol) glycerol, 5 mM beta-mercaptoethanol (BME), 0.1 % Tritin-X100, 2 µg/ml DNAse1, and protease inhibitors (84 µM leupeptin, 0.3 µM aprotinin, 1 µM pepstatin, 100 µM phenylmethylsulfonyl fluoride (PMSF)) and incubated with Ni-NTA agarose beads (Qiagen) for 2 hours. The bound protein was washed with 20 column volumes (CV) lysis buffer without lysozyme and protease inhibitors, 20 CV wash buffer (50 mM Tris, pH 8, 1 M NaCl, 20 mM Imidazole, 5% (wt/vol) glycerol, 5 mM beta-mercaptoethanol (BME), 0.009 % Tritin-X100) and eluted in four steps (1 CV per step) with wash buffer supplemented with increasing imidazole concentrations (50, 150, 250 and 400 mM). The eluted protein was dialyzed for 2h against gel filtration buffer (50 mM Tris, pH 8.0, 150 mM KCl, 1 mM DTT, 5% (wt/vol) glycerol). Ulp1 protease (0.75 mg per 30 ml) was added to the dialysis bag to remove the affinity tag and the dialysis reaction was performed overnight. The cleaved protein was incubated with 2 ml Ni-NTA agarose beads (Qiagen) to remove His_6_-Sumo and Ni-NTA agarose binding contaminants. The eluate was concentrated to 5 ml with a Viviaspin 20 (30.000 nominal molecular weight cut-off (MWCO), polyethersulfone (PES) membrane). Monomeric Chmp7 was isolated by gel filtration chromatography. Monomeric Chmp7 was concentrated to 24-42 µM using a Viviaspin 20 concentrator (30,000 nominal MWCO, PES membrane), aliquoted and snap-frozen in liquid N_2_. For binding experiments the protein was thawed on ice and cleared from aggregates by centrifugation for 20 min at 98,600 *g*, using a TLA-55 rotor (Beckman Coulter). Yields were 300-500 µg of protein per 1 L of bacterial culture.

*LEM2(NTD) purification:* 18 g of bacterial pellet were lysed (25 mM Tris, pH 7.0, 500 mM KCl, 10 mM Imidazole, 5 mM BME, 2 µg/ml DNAse1, and protease inhibitors) and incubated with Ni-NTA agarose beads (Qiagen) for 2 hours. The bound protein was washed with 40 CV lysis buffer without lysozyme and protease inhibitors and eluted in four steps (1 CV per step) with lysis buffer supplemented with increasing imidazole concentrations (50, 150, 250 and 400 mM). The eluted protein was dialyzed against SP column loading buffer (25 mM Tris, pH 7.0, 150 mM KCl, 1 mM DTT) and applied to a 5 ml HiTrap SP HP column (GE Healthcare), washed with loading buffer, and eluted with a gradient from 150 mM KCl to 500 mM KCl. The eluate was concentrated to 5 ml with a Viviaspin 20 (10,000 nominal MWCO, PES membrane) and monomeric His-Sumo-LEM2 (NTD) was isolated by gel filtration chromatography 25 mM Tris, pH 7.0, 150 mM KCl, 1 mM DTT). Monomeric His-Sumo-LEM2 (NTD) was concentrated to 20 µM and aliquots were snap-frozen in liquid N_2_. Yields were ~2.5 mg of protein per 1 L of bacterial culture. For binding experiments His-Sumo-LEM2 (NTD) was thawed on ice, spun for 10 min at 4°C at 16,627 *g*. The protein was used directly for binding or processed into untagged LEM2 (NTD). To obtain untagged protein 250 µg His-Sumo-LEM2 (NTD) were incubated with 30 µg Ulp1 for 1h at 4 °C and subsequently incubated with 100 µl Ni-NTA agarose beads (Qiagen) to remove cleaved His_6_-Sumo. The beads were removed by centrifugation and the supernatant was used for binding experiments.

*LEM2(CTD) purification:* Cells were suspended in buffer with 50mM Tris pH 8.0, 150mM KCl, 1mM DTT plus protease Inhibitor cocktail and were lysed by freeze-thaw cycles and the supernatant was harvested after 16,350 *g* for 45min. The lysate was incubated with Ni-beads (Qiagen) for 45min, washed with 20mM imidazole, and eluted with 750 mM imidazole. The eluted protein was dialyzed against gel-filtration buffer (50mM Tris pH 8.0, 150mM KCl, 1mM DTT) and applied to 120ml Hiload Superdex 75PG (GE healthcare). Fractions containing pure LEM2-CTD were collected and snap-frozen in liquid nitrogen. Yields were ~2.5 mg of protein per 1 L of bacterial culture.

### Protein binding experiments

Binding experiments were performed at room temperature using 40 µl Ni-NTA agarose beads (Qiagen) in 0.8 ml centrifuge columns (Pierce). Proteins were mixed and incubated for 1h. For control reactions proteins were incubated with corresponding buffers to mimic binding conditions. The protein reactions were added to the beads, which were equilibrated with binding buffer (25 mM, Tris pH 7.0, 150 mM KCl, 20 mM imidazole) and incubated for additional 45 min. The resin was washed with 20 CV of binding buffer. Excess His-Sumo-tagged-LEM2 (CTD or NTD) was eluted with three CV of binding buffer supplemented with 150 mM imidazole. Note that the majority of LEM2 and bound CHMP7 did not elute at this step. Protein remaining on beads was eluted with two CV 2x-SDS sample buffer and the eluate was analyzed with SDS-PAGE.

## Acknowledgements

We thank Dr. Brian Dalley and Dr. Tim Parnell for advice and expertise in whole genome sequencing and analysis, Dr. Janet Iwasa for graphic illustration, Dr. Mark Smith for assistance with confocal microscopy, Sarah M. Pick for technical assistance with protein purification, Dr. Doug Mackay for advice, Dr. Shelley Sazer for a NLS-GFP-LacZ nuclear integrity reporter, Dr. Yasushi Hiraoka for an Ish1-GFP strain, and Dr. John McCullough, Dr. Jeremy Carlton, and Dr. Patrick Lusk for stimulating conversations about unpublished results. Light and 2D transmission electron microscopy were performed in the Health Sciences Cores at the University of Utah. Microscopy equipment was obtained using a NCRR Shared Equipment Grant 1S10-RR024761-01. Our research was also supported by the Searle Scholars Program (A.F.), NIH 2P50-GM082545-06 (W.I.S., A.F., M.S.L., and P.J.B.), 1DP2-GM110772-01 (A.F.), 1R01-GM112080 (W.I.S.), the Huntsman Cancer Foundation and the Huntsman Cancer Institute Cancer Center Support Grant NIH P30CA042014 (K.S.U., W.I.S., A.F., the Genomics and Bioinformatics Shared Resource).

### Author Contributions

M.G., D.L., O.S.C., A.v.A., W.I.S., K.S.U., and A.F. designed research; M.G., D.L., O.S.C., A.v.A., M.S.L., M.J.R., and L.N. performed research; P.J.B. contributed new reagents/analytic tools; M.G., D.L., O.S.C., A.v.A., M.S.L., W.I.S., K.S.U., and A.F. analyzed data; and M.G., D.L., O.S.C., A.v.A., W.I.S., K.S.U., and A.F. wrote the paper.

### Conflict of Interest

The authors declare that they have no conflict of interest.

## Supplemental Supporting Information

### Supplemental Figures

**Figure S1.**
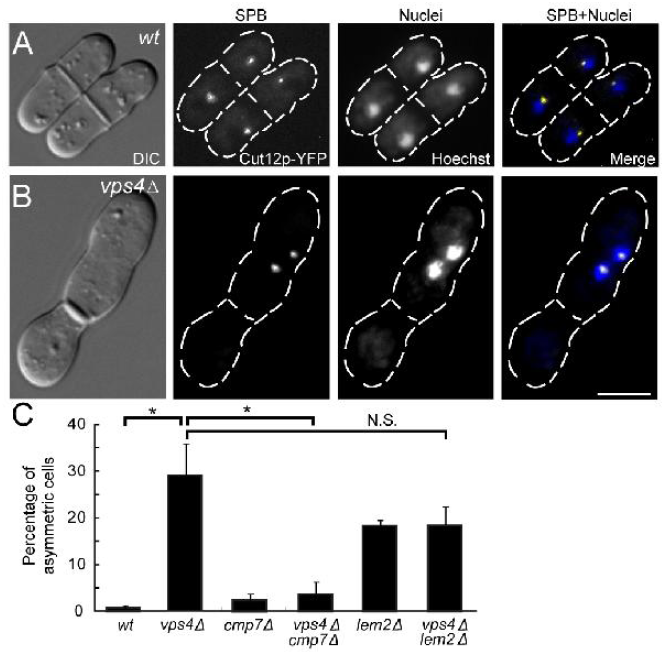
*vps4Δ* and *lem2Δ* exhibit asymmetric SPB segregation. (A) *wt* cells showing symmetric SPB segregation. (B) *vps4Δ* showing asymmetric SPB segregation. (C) Quantified percentages of asymmetric SPB segregation events in each genotype. Cells with septa were selected and scored for SPB separation patterns. Nuclei were stained with Hoechst. *wt*: 0.7±0.3%, n=103, 101, 101; *vps4Δ*: 29±7%, n=88, 103, 107; *cmp7Δ*: 2±1%, n=105, 103, 101; *vps4Δcmp7Δ:* 4±2%, n =117, 100, 104; *lem2Δ*: 18±1%, n= 102, 103, 100; *vps4Δlem2Δ*: 18±4%, n=102, 103, 101. (Scale bar: 5 μm).

**Figure S2.**
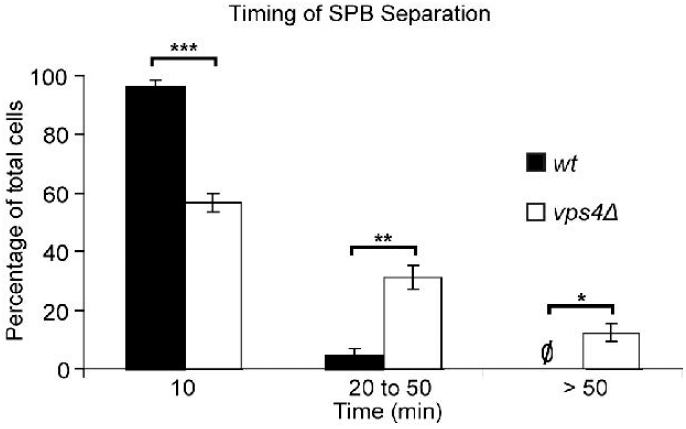
SPB separation is delayed in *vps4Δ* mutants. 16-hours time-lapse imaging (10 minute intervals) was conducted on *wt* and *vps4Δ* cells at 32°C, and the times required for SBP separation scored. Data were binned and plotted by mean frequency±SEM. *wt*: 10 minutes 96±2%; 20-50 minutes 4±2%; >50 minutes 0%; n=23, 22, 22; *vps4Δ*: 10 minutes 57±4%; 20-50 minutes 31±4%; >50 minutes 12±3%; n=30, 30, 30.

**Figure S3.**
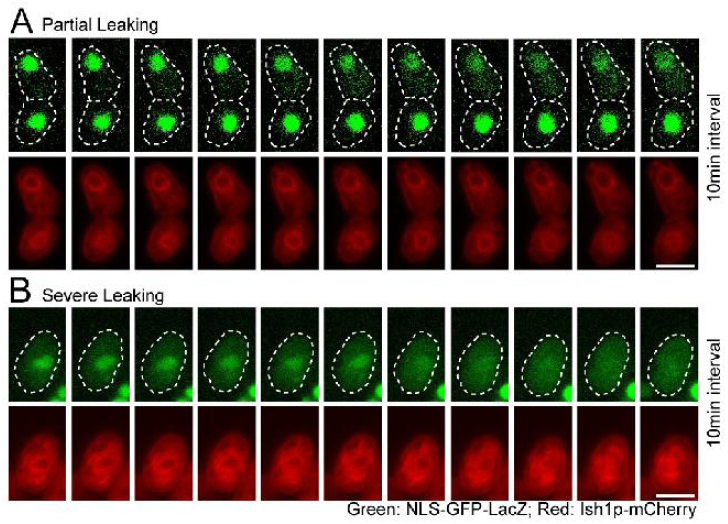
Time-lapse analysis of NLS-GFP-LacZ localization in *vps4Δ*. (A) Time-lapse images showing normal nuclear localization of NLS-GFP-LacZ converting to partial cytoplasmic localization. A pair of newly divided cells is shown over a time course of 100 minutes with 10-minute interval. (B) Time lapse images showing partial cytoplasmic localization of NLS-GFP-LacZ converting to a homogenous distribution throughout the cell. A single cell is shown over a time course of 100 minutes with 10-minute intervals. (Scale bars: 5 μm).

**Figure S4.**
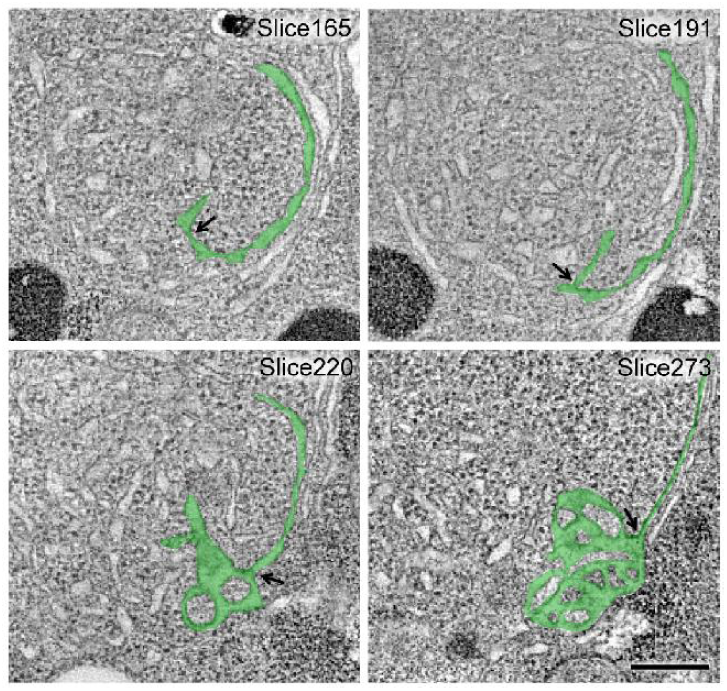
Karmellae and tubular extension are connected NE structures in *vps4Δ*. Selected tomographic slices of *vps4Δ* showing connected karmellae and tubular extensions. The relevant membranes are shaded light green and the junction is marked by black arrows. (Scale bars: 200 nm).

**Figure S5.**
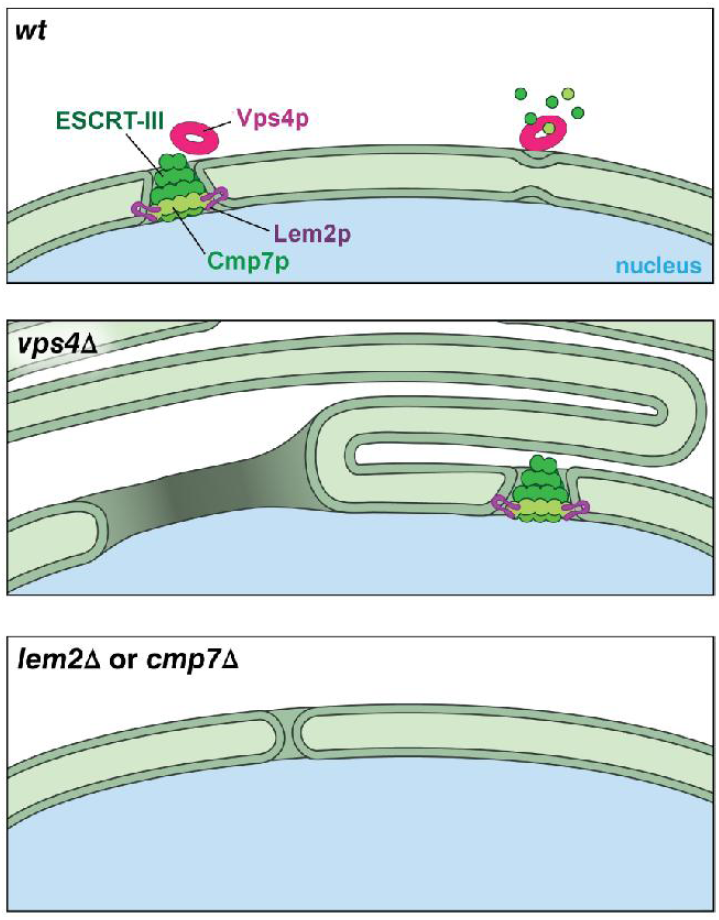
A cartoon model for the nuclear envelope phenotypes in *vps4Δ, lem2Δ, and cmp7Δ* in *S. pombe*. Top: in wild type, the exposed Lem2p recruits Cmp7p to initiate an ESCRT-III mediated nuclear envelope sealing event. That process is regulated and/or completed by the AAA ATPase, Vps4p. Middle: in *vps4Δ*, unrestricted Lem2p/Cmp7p activities induce the formation of karmellae and large gaps in nuclear envelope so that the proper progress of mitosis is disrupted. Bottom: In *lem2Δ* or *cmp7Δ*, mutant cells tolerate a limited number of small breaches and therefore show little or no growth defect.

**Figure S6.**
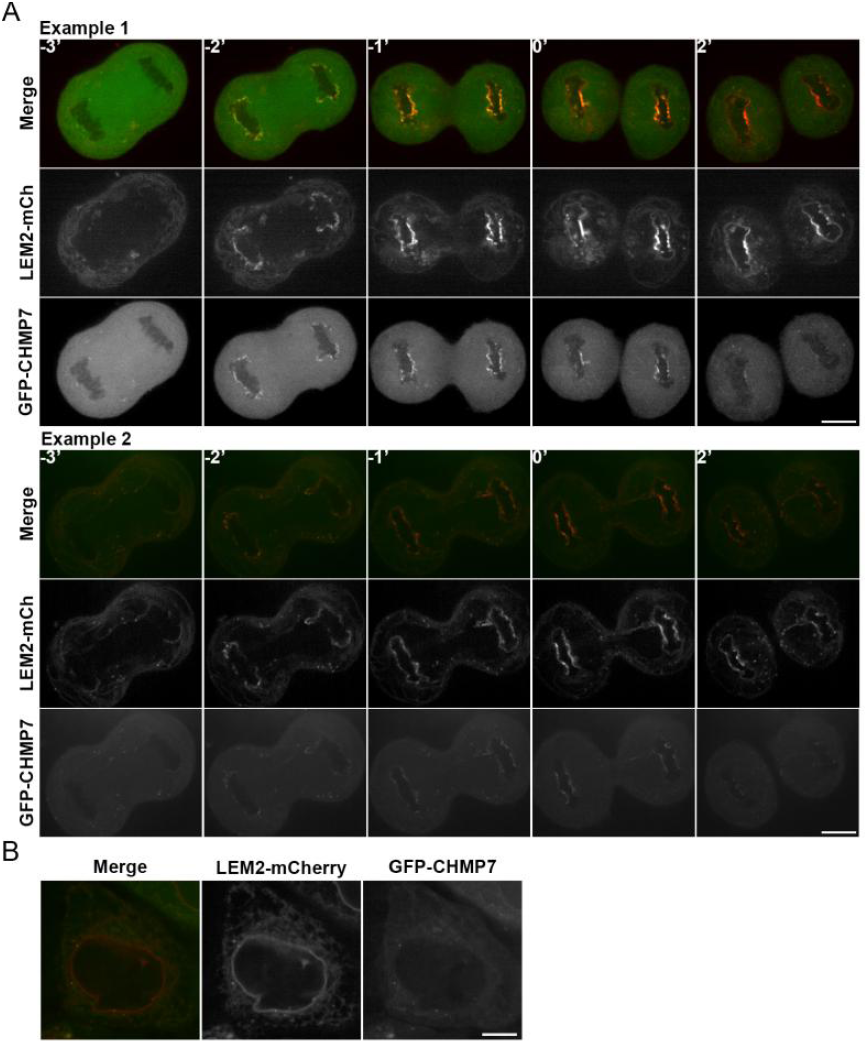
LEM2 and CHMP7 colocalization is restricted to a brief window specifically during anaphase. (A) Additional montages of cells expressing LEM2-mCherry and GFP-CHMP7 progressing through anaphase before complete furrow ingression (designated as t=0’). (B) Representative image of an interphase cell expressing LEM2-mCherry and GFP-CHMP7. (Scale bars: 10 μm).

**Figure S7.**
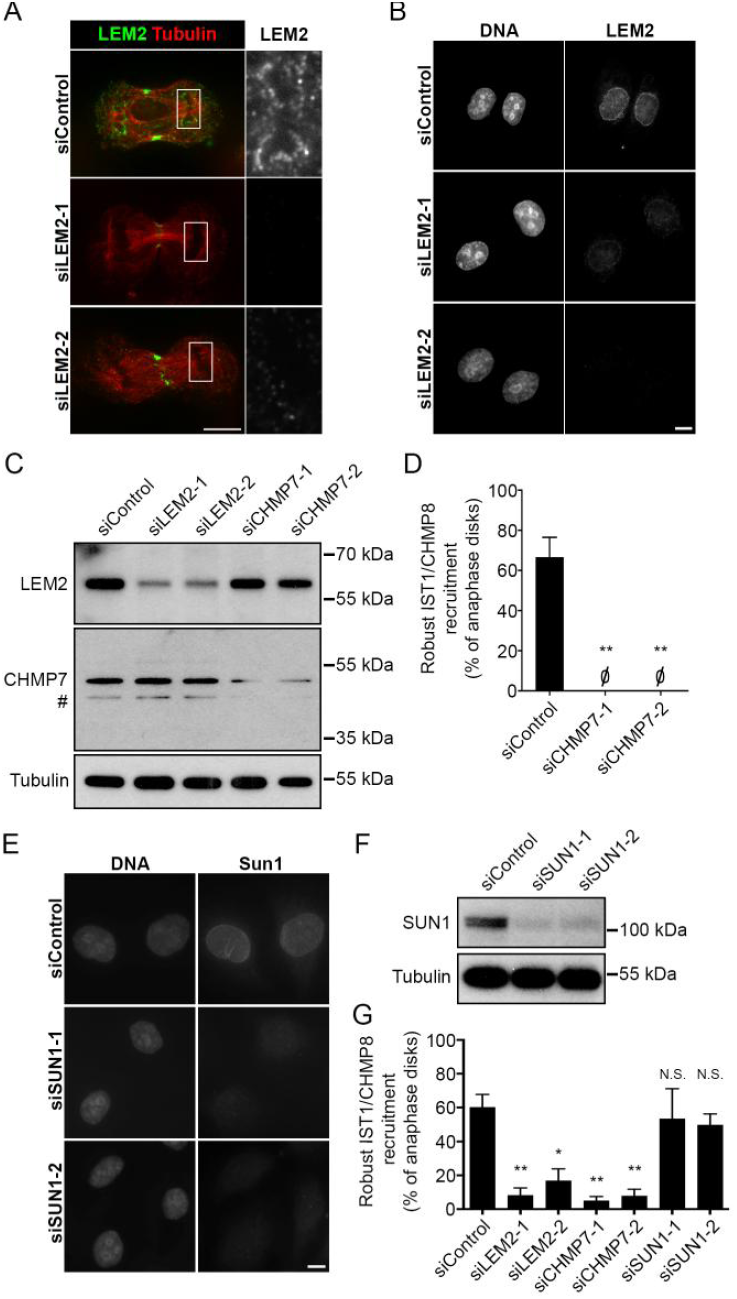
Control experiments for knockdown analysis. (A) Endogenous LEM2 detected at anaphase chromatin disks. Spinning disk confocal microscopy of representative anaphase cells 72 hours after treatment with siLEM2-1 or siLEM2-2 confirms that detection of signal at the chromatin surface is specific. Signal detected at the midzone with LEM2 antibody is non-specific, persisting after knockdown of LEM2 with two independent oligos. (B) Widefield microscopy of representative interphase cells comparing detection of endogenous LEM2 at the nuclear rim 72 hours after treatment with siControl or LEM2 specific oligos. (C) Specific depletion of LEM2 and CHMP7 with respective oligos used for RNAi is confirmed by immunoblot. # indicates a smaller protein product likely derived from CHMP7. (D) IST1/CHMP8 recruitment, assessed and graphed as described in Fig 7B, performed in parallel with LEM2 analysis in Fig 7F (siControl: 67±10%, n=104, 14, 40; siCHMP7-1: 0±0%, n=68, 26, 24; siCHMP7-2: 0±0%, n=24, 10, 48). (E) Widefield microscopy of representative interphase cells comparing detection of endogenous SUN1 at the nuclear rim 72 hours after treatment with siControl or SUN1-directed oligos. (F) Specific depletion of SUN1 by RNAi confirmed by immunoblot. (G) IST1/CHMP8 recruitment, assessed and graphed as described in Fig 7B, performed in parallel with CHMP2A analysis in Fig 7D (siControl: 60±8%, n=80, 42, 48; siLEM2-1: 8.2±4%, n=80, 82, 30; siLEM2-2: 17±7%, n=120, 82, 78; siCHMP7-1: 5±3%, n=76, 50, 58; siCHMP7-2: 8±4%, n=58, 46, 48; siSUN1A: 53±18%, n=58, 18, 20; siSUN1B: 50±7%, n=78, 44, 36). All graphs plot Mean±SEM.

**Figure S8.**
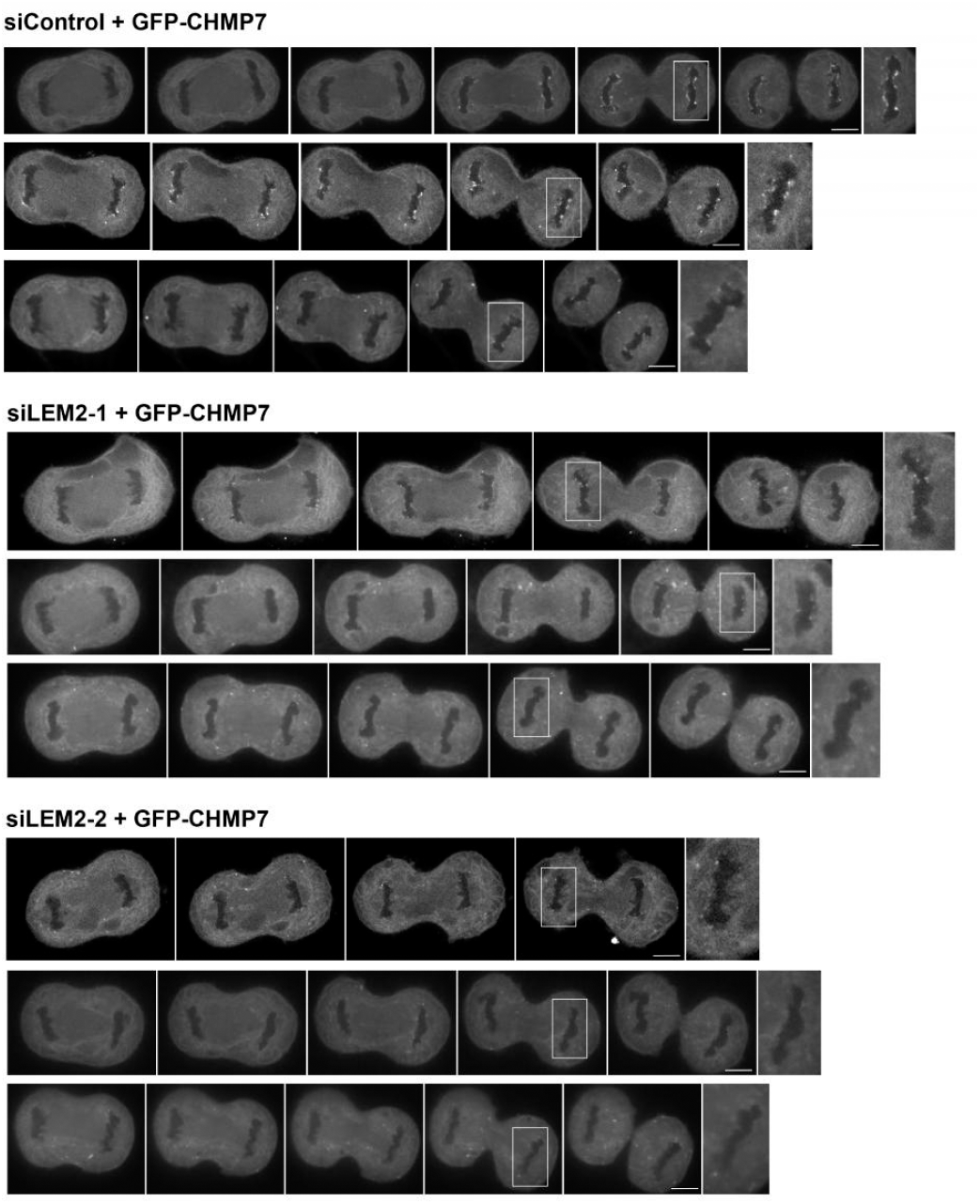
LEM2 plays a role in the recruitment of CHMP7 to anaphase chromatin disks. Additional montages of cells expressing GFP-CHMP7 and treated with siRNA, as indicated, progressing through anaphase.

**Figure S9 The C-terminal domain of LEM2 binds full-length CHMP7 *in vitro*.** (A) Protein domain structure of human LEM2. NTD, N-terminal domain; TM, transmembrane domain; CTD, C-terminal domain. (B) *In vitro* binding assay of full length human CHMP7 with immobilized human His-SUMO-LEM2(NTD) (AA 1-208, left) and (C) immobilized His-SUMO-LEM2(CTD) (AA 398-503, right). Samples are SDS-Buffer eluates analyzed by SDS-PAGE and Coomassie staining. Note that CHMP7 showed weak unspecific binding to the resin (B, lane 2) that was not increased by immobilized His-SUMO-LEM2(NTD) (B, lane 3). In contrast, immobilized His-SUMO-LEM2(CTD), increased CHMP7 binding significantly (C, compare lanes 3 and 4). Finally, LEM2 (NTD) did not bind immobilized LEM2 (CTD) (C, lane 5), and the presence of LEM2 (NTD) did not enhance CHMP7 binding to immobilized His-SUMO-LEM2 (CTD) (C, compare lanes 4 and 6).

### Movies

**Movie S1:** Segmented model of wild type NE from 150 tomographic slices. Color code: NE‐‐‐ green; NPCs‐‐‐red.

**Movie S2:** Segmented model of *vps4Δ* NE from 100 tomographic slices. Color code: NE‐‐‐ green; NPCs‐‐‐red; Karmellae‐‐‐gold; A whorl of tubules‐‐‐purple.

**Movie S3:** Segmented model of *vps4Δ* NE from 200 tomographic slices. Color code: NE‐‐‐ green; NPCs‐‐‐red.

**Movie S4:** Segmented model of another *vps4Δ* NE from 150 tomographic slices. Color code: NE‐‐‐green; NPCs‐‐‐red; SPB‐‐‐yellow.

**Movie S5:** Movie of a cell expressing GFP-CHMP7 and LEM2-mCherry progressing through anaphase. The interval between each frame is 20 seconds. Images from this movie are shown as a montage in Fig 6A, using 1 minute intervals.

**Movie S6:** Movie of a cell treated with siControl expressing GFP-CHMP7 and H2B-mCherry progressing through anaphase. The interval between each frame is 30 seconds.

**Movie S7:** Movie of a cell treated with siLEM2-1 expressing GFP-CHMP7 and H2B-mCherry progressing through anaphase. The interval between each frame is 30 seconds.

**Movie S8:** A series of Z-stack images of an illustrative anaphase B cell treated with siControl. IST1/CHMP8 is shown in green and tubulin in red.

**Table S1:** Spontaneous suppressor alleles of *vps4Δ* identified in this study.

| Suppressor | Suppressor Mutation Alleles | Mutation | Total Number Valid Reads | Number of Supporting Reads | Percentage of Supporting Reads | Residues Affected | Length of Peptide Translated |
| --- | --- | --- | --- | --- | --- | --- | --- |
| *A. cmp7*<sup>†</sup> |  |  |  |  |  |  |  |
| 1 | *cmp7-1* | Frameshift | 26 | 25 | 96 % | 24 - 449 | 24 |
| 2 | *cmp7-2* | Nonsense | 35 | 35 | 100 % | W49*(stop) | 48 |
| 3 | *cmp7-3* | Nonsense | 20 | 19 | 95 % | Q139*(stop) | 138 |
| 4 | *cmp7-4* | Frameshift | 15 | 15 | 100 % | 280 - 449 | 284 |
| 5 | *cmp7-5* | Frameshift | 10 | 10 | 100 % | 126 - 449 | 155 |
| 6 | *cmp7-6* | Frameshift | 31 | 31 | 100 % | 115 - 449 | 148 |
| 7 | *cmp7-7* | Frameshift | 19 | 19 | 100 % | 285 - 449 | 287 |
| *B. lem2*<sup>‡</sup> |  |  |  |  |  |  |  |
| 8 | *lem2-1*<sup>§</sup> | Frameshift | 20 | 20 | 100 % | 550 – 668 | 561 |
| 9 | *lem2-1* | Frameshift | 17 | 17 | 100 % | 550 – 668 | 561 |
| 10 | *lem2-1* | Frameshift | 23 | 23 | 100 % | 550 – 668 | 561 |
| 11 | *lem2-1* | Frameshift | 22 | 22 | 100 % | 550 – 668 | 561 |
| 12 | *lem2-1* | Frameshift | 24 | 23 | 95 % | 550 – 668 | 561 |

\* Whole genome sequencing of *wild type* (n=2), *vps4* mutant (n=2) and spontaneous suppressors (n=12) uncovered mutant alleles responsible for rescuing growth defects of *vps4*.
<sup>†</sup> Wild type *cmp7* encodes a peptide of 449 amino acids.
<sup>‡</sup> Wild type *lem2* encodes a peptide of 668 amino acids.
<sup>§</sup> The *lem2-1* mutant allele had a single T inserted within a series of 7 T residues corresponding to nucleotides 1643 – 1649 (relative to the *lem2* ATG).

**Table S2:** Quantification of IST1/CHMP8 recruitment in mammalian cells.

| IST1 |  | Figure 7B |  |  |  | Figure S6G |  |  |  | Figure S7D |  |  |  |
| --- | --- | --- | --- | --- | --- | --- | --- | --- | --- | --- | --- | --- | --- |
|  |  | Exp 1 | Exp 2 | Exp 3 | Mean | Exp 4 | Exp 5 | Exp 6 | Mean | Exp 7 | Exp 8 | Exp 9 | Mean |
| siControl | Robust* | 56 | 76 | 58 | **63** | 45 | 67 | 69 | **60** | 81 | 71 | 48 | **67** |
|  | Weak | 39 | 19 | 38 | **32** | 40 | 26 | 25 | **30** | 19 | 29 | 45 | **31** |
|  | Absent | 5 | 5 | 4 | **5** | 15 | 7 | 6 | **9** | 0 | 0 | 7 | **2** |
| siLEM2-1 | Robust | 0 | 0 | 0 | **0** | 0 | 15 | 10 | **8** |  |  |  |  |
|  | Weak | 53 | 52 | 17 | **41** | 60 | 39 | 47 | **49** |  |  |  |  |
|  | Absent | 47 | 48 | 83 | **59** | 40 | 46 | 43 | **43** |  |  |  |  |
| siLEM2-2 | Robust | 0 | 18 | 0 | **6** | 5 | 16 | 29 | **17** |  |  |  |  |
|  | Weak | 32 | 50 | 100 | **61** | 34 | 46 | 35 | **38** |  |  |  |  |
|  | Absent | 68 | 32 | 0 | **33** | 61 | 38 | 36 | **45** |  |  |  |  |
| siCHMP7-1 | Robust | 0 | 0 | 0 | **0** | 0 | 8 | 7 | **5** | 0 | 0 | 0 | **0** |
|  | Weak | 2 | 0 | 0 | **1** | 8 | 22 | 29 | **20** | 3 | 4 | 0 | **2** |
|  | Absent | 98 | 100 | 100 | **99** | 92 | 70 | 64 | **75** | 97 | 96 | 100 | **98** |
| siCHMP7-2 | Robust | 0 | 0 | 0 | **0** | 0 | 13 | 10 | **8** | 0 | 0 | 0 | **0** |
|  | Weak | 9 | 0 | 0 | **3** | 10 | 50 | 50 | **37** | 0 | 0 | 0 | **0** |
|  | Absent | 91 | 100 | 100 | **97** | 90 | 37 | 40 | **56** | 100 | 100 | 100 | **100** |
| siSun1A | Robust |  |  |  |  | 36 | 89 | 35 | **53** |  |  |  |  |
|  | Weak |  |  |  |  | 55 | 11 | 60 | **42** |  |  |  |  |
|  | Absent |  |  |  |  | 9 | 0 | 5 | **5** |  |  |  |  |
| siSun1B | Robust |  |  |  |  | 49 | 61 | 39 | **50** |  |  |  |  |
|  | Weak |  |  |  |  | 45 | 32 | 56 | **44** |  |  |  |  |
|  | Absent |  |  |  |  | 6 | 7 | 6 | **6** |  |  |  |  |

**Table S3:** Quantification of CHMP2A recruitment in mammalian cells.

| CHMP2A |  | Figure 7D |  |  |  |
| --- | --- | --- | --- | --- | --- |
|  |  | Exp 4 | Exp 5 | Exp 6 | Mean |
| siControl | Robust | 73 | 83 | 54 | **70** |
|  | Weak | 23 | 17 | 38 | **26** |
|  | Absent | 4 | 0 | 8 | **4** |
| siLEM2-1 | Robust | 12 | 16 | 0 | **9** |
|  | Weak | 70 | 63 | 61 | **65** |
|  | Absent | 18 | 21 | 39 | **26** |
| siLEM2-2 | Robust | 19 | 4 | 10 | **11** |
|  | Weak | 64 | 50 | 52 | **55** |
|  | Absent | 16 | 46 | 38 | **33** |
| siCHMP7-1 | Robust | 6 | 5 | 2 | **4** |
|  | Weak | 26 | 38 | 53 | **39** |
|  | Absent | 68 | 57 | 45 | **57** |
| siCHMP7-2 | Robust | 5 | 28 | 30 | **21** |
|  | Weak | 38 | 60 | 57 | **52** |
|  | Absent | 56 | 12 | 13 | **27** |

**Table S4:** Quantification of LEM2 recruitment in mammalian cells.

| LEM2 |  | Figure 7F |  |  |  |
| --- | --- | --- | --- | --- | --- |
|  |  | Exp 7 | Exp 8 | Exp 9 | Mean |
| siControl | Present | 100 | 97 | 87 | **95** |
|  | Absent | 0 | 3 | 13 | **5** |
| siCHMP7-1 | Present | 100 | 81 | 87 | **89** |
|  | Absent | 0 | 19 | 13 | **11** |
| siCHMP7-2 | Present | 100 | 96 | 100 | **99** |
|  | Absent | 0 | 4 | 0 | **1** |

\*Numbers reported are the percentage of chromatin disks scored as robust, weak or absent for IST1, CHMP2A, and LEM2 for each experiment.

**Table S5:**
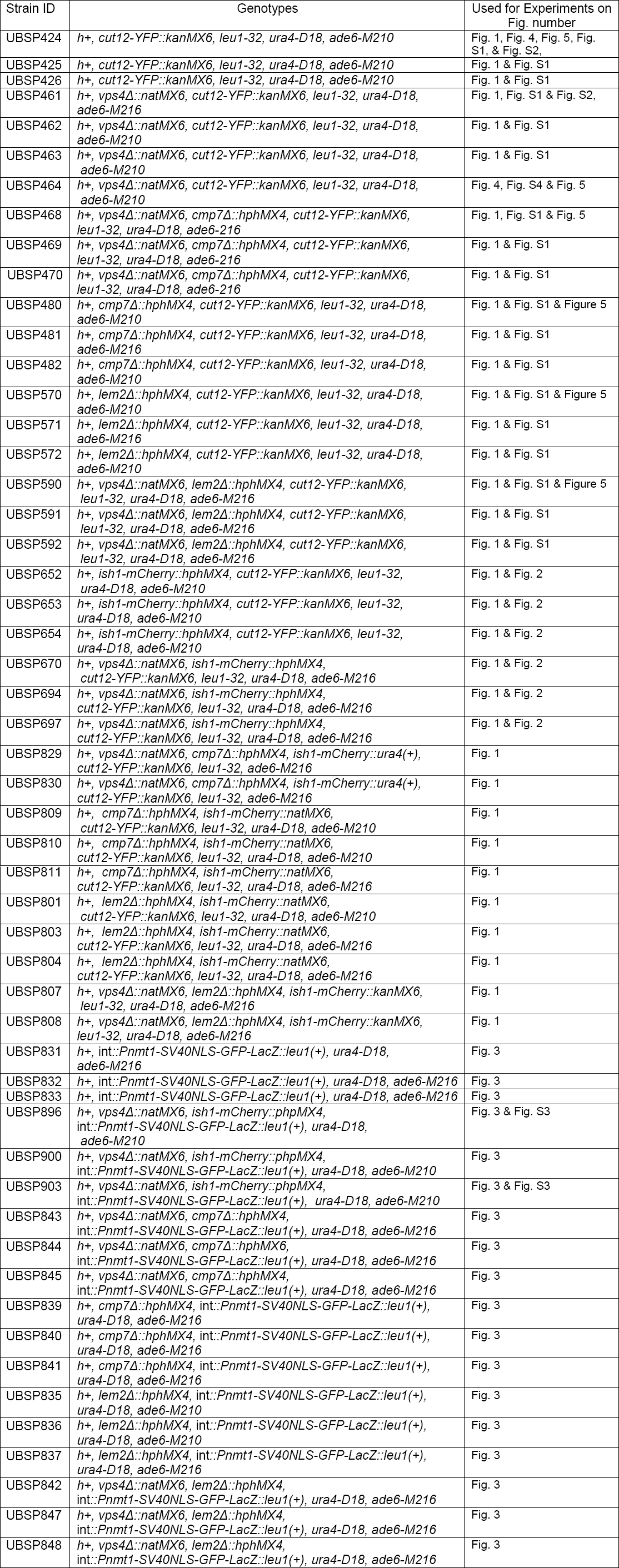
Yeast strains constructed for this study.

